# Grapevine Red Blotch Virus Induces Haplotype-specific Genetic and Epigenetic Responses

**DOI:** 10.64898/2026.01.29.702617

**Authors:** Christian Mandelli, Samuel C. Talbot, Ahmed Z. Alsulaimawi, Laurent G. Deluc

## Abstract

Haplotype-resolved genomes provide a powerful framework for uncovering allele-specific regulatory mechanisms obscured in collapsed diploid references. Despite growing evidence of cultivar- and clone-dependent variation in disease severity caused by Grapevine Red Blotch Virus (GRBV), the contribution of haplotype-specific transcriptional and epigenetic regulation to host–virus interactions remains poorly understood. Here, we integrate a haplotype-resolved grapevine genome with time-resolved transcriptomic and whole-genome bisulfite sequencing to dissect allele-specific regulatory responses to GRBV infection. Allele-aware RNA-seq analysis revealed extensive haplotype-dependent transcriptional remodeling, with both shared and divergent temporal expression programs across infection stages. Soft clustering and weighted gene co-expression network analysis (WGCNA) identified haplotype-specific co-expression modules and regulatory hubs, uncovering asymmetric network organization and distinct antiviral strategies between parental haplotypes. Notably, chloroplast-associated defense networks emerged as conserved but highly infection-sensitive modules, exhibiting haplotype-dependent recovery or sustained disruption during infection. Methylome profiling demonstrated that GRBV infection induces pronounced haplotype-specific epigenetic reprogramming, with differential DNA methylation concentrated in promoter-proximal and transposable element–associated regions. Together, our results demonstrate that haplotype-resolved, multi-omic analyses reveal regulatory complexity and divergent antiviral strategies that are hidden by collapsed-genome approaches. This work provides new mechanistic insight into grapevine–virus interactions and lays a foundation for leveraging allelic variation to improve disease resilience in clonally propagated crops.

## Introduction

Haplotype-resolved genomes can reveal allelic diversity, regulatory asymmetries, and stress-response mechanisms that remain hidden in collapsed assemblies. In wine grape (*Vitis vinifera*), which is highly heterozygous (Xiao et al. 2025) and exhibits cultivar-specific disease responses, haplotype-level resolution is essential for understanding how distinct alleles influence host–pathogen interactions.

Upon domestication and clonal propagation, *Vitis vinifera* has accumulated extensive genetic diversity. This variation serves as a critical resource for breeding and cultivar improvement, yet many genes underlying key agronomic traits remain poorly characterized.

Advances in high-throughput sequencing have accelerated the discovery of indels, SNPs, and structural variants (SVs), thereby refining our understanding of the architecture of the grapevine genome. Recent chromosome-scale assemblies (Wang et al. 2024) have revealed the extent of genomic variation across grape accessions.

Importantly, haplotype-resolved assemblies have proven essential for linking allelic variants to phenotypes, enabling the identification of genes involved in berry pigmentation, stress responses, clonal divergence, and other agronomic traits (Calderón et al. 2024; Cochetel et al. 2023; Shirasawa et al. 2022; Wang et al. 2024; Zhang et al. 2023). These studies collectively demonstrate that haplotype-resolved genomic analyses can reveal functional diversity hidden in collapsed diploid references, providing a framework for dissecting allele-specific contributions to trait expression.

Despite these advances, most functional genomics studies in grapevine still rely on collapsed diploid references (Zhao et al. 2025) or unphased gene models, which hide the allele-specific regulatory architecture that can influence transcriptional output (Callipo et al. 2025; Cardone et al. 2016; Martínez-Zapater et al. 2010). In highly heterozygous perennial crops, alleles frequently differ not only in coding sequence but also in regulatory potential, giving rise to allele-specific expression (ASE) and asymmetric transcriptional responses across haplotypes (Wong et al. 2018). This regulatory heterogeneity is manifested at the phenotypic level by the coexistence of resistant and susceptible cultivars (Trapp et al. 2025) in response to abiotic or biotic stress. These differences in stress response might arise from how alleles are regulated or networked rather than from gene presence alone. Consequently, without haplotype-resolved references, such regulatory asymmetries are difficult to detect, and expression signals from distinct alleles may be underestimated or completely undiscovered.

This limitation is particularly consequential in the context of biotic stress responses, where fine-scale regulatory variation can determine the timing, magnitude, and tissue specificity of defense gene activation. Plant–pathogen interactions often involve rapid transcriptional reprogramming, mediated by resistance (R) genes, signaling components, and downstream defense pathways. In grapevine, disease outcomes frequently vary among cultivars and even among clonal lineages (Boso et al. 2004; Callipo et al. 2025; Murolo and Romanazzi 2014), suggesting that allelic differences might contribute to pathogen susceptibility or tolerance. However, the extent to which individual haplotypes differentially contribute to defense signaling and pathogen response remains largely unexplored.

To date, haplotype-resolved genomic approaches have been only sparsely applied to grapevine–pathogen interaction studies (Guo et al. 2025; Massonnet et al. 2022). The potential of phased assemblies to elucidate host–pathogen dynamics has not been systematically leveraged. There is a lack of integrative analyses that combine haplotype-resolved genomes with transcriptomic or network-based approaches to disentangle allele-specific regulation during infection. Addressing this gap is essential for moving beyond descriptive variation toward a more quantitative understanding of how allelic diversity shapes grapevine immune responses, with direct implications for disease management, breeding, and the durability of resistance traits in clonally propagated cultivars.

Grapevine Red Blotch Virus (GRBV) is an emerging, economically significant DNA virus associated with Grapevine Red Blotch Disease, which causes delayed ripening, reduced sugar accumulation, altered secondary metabolism, and substantial quality losses in wine grape cultivars. Since its identification (Al Rwahnih et al. 2013), research on GRBV has primarily focused on symptomatology, epidemiology, vector transmission, and whole-plant physiological effects (Cauduro Girardello et al. 2020; Copp et al. 2025; Flasco et al. 2024, 2023; Setiono et al. 2018), with transcriptomic studies conducted using collapsed diploid references and bulk gene-level analyses (Ault et al. 2025; Blanco-Ulate et al. 2017; Rumbaugh et al. 2022). Despite pronounced cultivar- and clone-dependent variability in disease severity and phenotypic outcomes (Rumbaugh et al. 2021; Wallis and Sudarshana 2016), little is known about how allelic diversity and haplotype-specific regulatory programs shape grapevine responses to GRBV infection.

In this study, we address this gap by integrating a haplotype-resolved grapevine genome with time-resolved transcriptomic profiling of GRBV-infected tissues to dissect allele-specific regulatory responses at both the gene and network levels.

By combining allele-aware RNA-seq quantification with clustering and network approaches, including soft clustering and weighted gene co-expression network analysis (WGCNA), we characterize dynamic transcriptional trajectories across infection stages and identify haplotype-specific modules associated with the viral response. This integrative framework allowed us to disentangle shared versus haplotype-biased regulatory programs, reveal asymmetric network organization between alleles, and identify candidate hub genes that may contribute to the GRBV response. Together, our results demonstrate that haplotype-resolved, network-based analyses uncover regulatory complexity that is otherwise obscured in collapsed references, providing new mechanistic insights into grapevine–virus interactions and establishing a foundation for leveraging allelic variation in disease management and breeding strategies.

## Materials and Methods

### Plant Material

A population of microvine V4 plantlets (04C023V0004), derived from a cross between Grenache and L1 Pinot Meunier GAI1 mutant cultivars (Chaïb et al. 2010), was cultured for eight weeks in tissue culture on Murashige and Skoog (MS) basal rooting medium (Murashige and Skoog 1962), optimized for grapevine tissue culture (Kurth et al. 2012). The sterilized medium was poured into a double-Magenta box system. The plantlets were maintained at 25°C under a 16/8-hour photoperiod with a light intensity of 100 µmol m⁻² s⁻¹.

### GRBV artificial infection of microvine plants via vacuum-assisted *Agrobacterium tumefaciens-mediated* infiltration

A 100 mL MG/L liquid medium (Garfinkel and Nester, 1980), supplemented with 50 mg/L kanamycin and 10 mg/L rifampicin, was inoculated with a single colony of *Agrobacterium tumefaciens* strain EHA105 carrying a bitmer construct (Yepes et al., 2018) with the infectious GRBV NY358 clone (clade 2, GenBank JQ901105.2; Krenz et al., 2012). The culture was incubated at 29°C in a shaker incubator at 225 RPM for 36 hours. Upon reaching an OD600nm of 2.0, the culture was divided into 50 mL tubes and centrifuged at 1600 g for 5 minutes. The supernatant was discarded, and the bacterial pellets were washed twice with 5 mL of induction medium (30 mM MES, 1.7 mM NaH2PO4, 1% mannitol, pH 5.5 adjusted with 1M KOH) containing 200 μM acetosyringone at 1600 g for 5 minutes, as previously reported (Vaghchhipawala et al., 2011). The pellets were then resuspended in 400 mL of induction medium to achieve an OD600nm of 1.0 and transferred to a sterile 500 mL Erlenmeyer flask. This suspension was incubated at room temperature (22°C ± 2) with gentle shaking (<100 RPM) for 8 hours. After incubation, the induced bacteria were pelleted and resuspended in infiltration medium (Vaghchhipawala et al., 2011) to an OD600nm of 0.5, yielding 1.5 L of infiltration solution.

Primary and secondary roots of 8-week-old V4 microvine plants (five biological replicates per time point) were trimmed by one-third. Each plant’s roots were placed in the infiltration solution at the bottom of a vacuum chamber. Control plants (five biological replicates per time point) underwent the same procedure but with induced bacteria lacking the GRBV construct. Two vacuum cycles (650 mmHg or 0.855 atm) of 2 minutes each, with a 30-second release between cycles, were applied using a pressure-control valve system (Noshok Inc., OH, USA). Post-infiltration, plants were transferred to double-Magenta boxes with sterile vermiculite and 50 mL of the growth medium described in the Plant Material section for recovery. At 3 days post-infiltration (3 dpi, T1), an additional 50 mL of growth medium containing 1 g/L ticarcillin and clavulanic acid was added to eliminate residual bacteria.

### Total RNA sequencing

Leaves, petioles, and stems from infiltrated plants were sampled at three-, six-, and 12-dpi (T1 through T3, respectively). Samples from each plant were then pooled and ground into a fine powder using liquid nitrogen. For each biological replicate, 100 mg of frozen tissue was used for total RNA extraction with the RNeasy Plant Kit (Qiagen, Hilden, Germany) following a modified protocol (Mandelli and Deluc 2025). On-column DNase treatment was performed by incubating samples with approximately 6 Kunitz units of TURBO™ DNase (Invitrogen™, Thermo Fisher Scientific, Waltham, USA) at 37°C for 30 minutes. Three washes with RPE buffer were conducted to remove residual salts from the binding column. RNA was eluted using nuclease-free water (Qiagen, Hilden, Germany) pre-warmed to 65°C. To check for genomic DNA (gDNA) contamination, 250 ng of extracted RNA was used as a template for PCR targeting the microvine housekeeping gene VitviActin1 (Vitvi04g01613) (Rossdeutsch et al. 2016). The concentration and integrity of total RNA samples were evaluated at the Center for Quantitative Life Sciences (CQLS, Oregon State University, Corvallis, USA) using the Agilent Bioanalyzer 2100 (Agilent Technologies, Santa Clara, USA) for integrity and the Qubit fluorometer (Thermo Fisher Scientific, Waltham, USA) for quantification. Library preparation for RNA sequencing was performed using the NEBNext Ultra II kit (E7760S; NEB) and the NEBNext Poly(A) mRNA Magnetic Isolation Module (E7490S; NEB) according to the manufacturer’s instructions.

Library quality and size distribution were assessed using the Agilent Bioanalyzer 2100 with a High Sensitivity DNA Chip (Agilent Technologies, Santa Clara, USA). Library quantification was performed using the Qubit fluorometer (Thermo Fisher Scientific, Waltham, MA, USA) and validated by quantitative PCR (qPCR) to ensure accurate molarity for sequencing. The prepared libraries were pooled in equimolar ratios and submitted for sequencing on an Illumina NextSeq 2000 platform using a paired-end 100-nt mode, at the Center for Quantitative Life Sciences (CQLS, Oregon State University, Corvallis, USA).

### Whole Genome Bisulfite Sequencing

Genomic DNA was isolated using the DNeasy Plant Mini Kit (QIAGEN, Hilden, Germany), following a modified protocol (Mandelli and Deluc 2025). For this procedure, infected tissues (leaves, stems, and petioles) were sampled at 12- and 24-dpi (three biological replicates per collection time). For library preparation, the NEBNext Enzymatic Methyl-seq Kit (E7120S; NEB) was used according to the manufacturer’s instructions. Library quality and size distribution were assessed at the Center for Quantitative Life Sciences (CQLS, Oregon State University, Corvallis, USA) using the Agilent Bioanalyzer 2100 with a High Sensitivity DNA Chip (Agilent Technologies, Santa Clara, USA). Library quantification was performed with the Qubit dsDNA HS Assay Kit (Thermo Fisher Scientific, Waltham, USA) and validated by qPCR. Sequencing was conducted on an Illumina NextSeq 2000 platform with a single-end 50-nt mode, incorporating 5% PhiX as recommended by NEB.

### Bioinformatic Analysis

Bioinformatic analyses of the sequencing data were performed using standardized pipelines tailored to each sequencing experiment. For RNA sequencing, the nf-core/rnaseq pipeline (version 3.14.0) (Ewels et al. 2020) was utilized with Salmon (version 1.10.1) as the aligner for transcript quantification. Quality control was conducted using MultiQC (version 1.14) to generate comprehensive reports on sequencing metrics. Differential expression analysis was subsequently performed in R (version 4.3.2) using DESeq2 (version 1.38.0) to identify differentially expressed genes across the sampled time points and between infected and control tissues.

For whole-genome bisulfite sequencing (WGBS), the nf-core/methylseq pipeline (version 2.6.0) (Ewels et al. 2020) was used with the Bismark workflow (version 0.24.2) for alignment to the reference genome and methylation calling. Downstream analysis in R was conducted using the MethylKit package (Akalin et al. 2012) to map methylation levels, categorize methylation sites into genes, promoter regions, or intergenic regions, and assign them to specific chromosomes. For the RNA sequencing and WGBS data analysis described above, the haplotype-resolved genome assemblies and annotations of the microvine V4 (04C023V0004) were used.

### Time-series clustering and functional enrichment

Temporal co-expression analysis was performed using Mfuzz (v2.60.0) (Kumar and E. Futschik 2007) in R (v4.3.2) to identify coordinated expression trends among infection-responsive DEGs. For each haplotype, variance-stabilized counts from DESeq2 were averaged across infected biological replicates at each time point (T1–T3) and standardized by gene. DEGs detected at any infection stage (adjusted p < 0.05) were combined to define the clustering input. Genes with low temporal variance (standard deviation < 0.05) were excluded prior to analysis. The optimal fuzzification parameter m was estimated using the mestimate() function, and soft clustering was performed with k = 6 and k = 10 clusters. Minimum membership values (0–1) quantified the degree of association of each gene to its assigned temporal trend; genes with membership ≥ 0.50 were considered high-confidence members. Cluster-specific expression profiles were visualized both as mean trajectories and individual gene “spaghetti” plots.

To interpret biological processes underlying each temporal pattern, GO enrichment analyses were performed for all and high-confidence cluster members using topGO (v2.54.0) (Alexa and Rahnenfuhrer n.d.) with weight01 Fisher’s exact tests, based on Blast2GO-derived gene-to-GO mappings. Significant terms (Benjamini–Hochberg adjusted p < 0.05) were summarized across Biological Process (BP), Molecular Function (MF), and Cellular Component (CC) ontologies, and visualized as composite dot plots of representative enriched functions per cluster.

### Cross-haplotype comparison of co-expression dynamics

To assess conservation of temporal expression patterns between haplotypes, Mfuzz cluster assignments for each haplotype were compared using a reciprocal best-hit gene-pairing matrix. Orthologous transcript pairs were identified from reciprocal BLAST results, and cluster-level overlap was quantified as the percentage of paired genes assigned to equivalent expression trends in both haplotypes. Pairing matrices were visualized as bidirectional heatmaps, and summary bar plots were generated to show the proportions of shared and haplotype-specific genes across clustering resolutions (k = 6–12) and membership thresholds (≥ 0.10–1.00).

### WGCNA Analysis of RNA-seq Data

Weighted gene co-expression network analysis (WGCNA) (Langfelder and Horvath 2008) was performed independently for each haplotype to identify modules of co-expressed genes and to characterize their relationships with infection-associated traits. Only genes that were expressed (normalized counts > 1 in at least 80% of samples) were retained for network construction. Prior to analysis, variance-stabilized expression values from the DESeq2 pipeline were used as input to minimize heteroscedasticity across samples.

For each haplotype-specific dataset, the pairwise Pearson correlations between all expressed alleles were computed and transformed into an adjacency matrix using a signed power function. The soft-thresholding power (β) was selected as the lowest value at which the scale-free topology fit index (R²) first exceeded 0.8, ensuring approximate scale-free network topology. The adjacency matrix was then converted into a topological overlap matrix (TOM).

Modules of highly co-expressed genes were identified using the blockwiseModules() function in WGCNA, with the following parameters: TOMType = “unsigned”, networkType = “signed”, deepSplit = 3, minModuleSize = 250, and mergeCutHeight = 0.25. Modules whose eigengenes exhibited Pearson correlation greater than 0.75 were merged to minimize redundancy. Each module was assigned a unique color label for visualization.

To identify biologically relevant modules, module eigengenes were correlated with infection traits. Both Pearson and biweight midcorrelation (bicor) methods were used to ensure robustness to outliers. Modules showing significant associations (|r| > 0.4, p < 0.05) with infection traits were considered infection responsive. Within these modules, hub genes were defined as those showing high module membership (kME > 0.8) and strong gene–trait correlations (|r| > 0.5).

To assess the conservation of co-expression structures between haplotypes, module preservation analysis was performed using the modulePreservation() function on matched expression matrices. The analysis quantified the extent to which modules defined in one haplotype were preserved in the other, based on the Zsummary statistic (Zsummary > 10 = strongly preserved; 2 < Zsummary < 10 = moderately preserved). Modules showing low preservation were considered haplotype-specific and were further explored for unique biological functions.

For functional characterization, GO enrichment analysis was performed on each module using topGO with the “weight01” algorithm and Fisher’s exact test, using Blast2GO-derived annotations as the reference background. Enriched GO terms (FDR < 0.05) were summarized and visualized to highlight major biological processes represented within each module. Network edges corresponding to high TOM similarity (TOM ≥ 0.1) were exported to Cytoscape (v3.9.1) using the exportNetworkToCytoscape() function for hub gene visualization and network topology inspection.

### qPCR Experiments

To assess GRBV RepA and CP gene expression dynamics, the total RNA (500 ng) extracted from samples collected at zero-, three-, six-, and 12-dpi (Total RNA Sequencing section) was reverse transcribed. The expression levels of key GRBV genes (RepA or and CP) were quantified using real-time PCR. The grape VitviActin1 (Vitvi04g01613) gene served as the housekeeping gene for normalization (Rossdeutsch et al. 2016). Transcript abundance of the GRBV genes was measured using specific primers (Supplementary Table S1). qPCR reactions consisted of 40 cycles of two-step cycling at 95°C for 15 s and 60°C for 1 min. Gene expression was calculated using the ΔΔCt method with VitviActin1 as the reference gene (Livak and Schmittgen 2001).

### Data and code availability

The raw sequencing data generated in this study have been deposited in the NCBI Sequence Read Archive (SRA) under the BioProject accession number PRJNA1407057. Processed data files, including haplotype-aware expression matrices, methylation summaries, and co-expression network outputs, are available from the corresponding author upon reasonable request.

All custom scripts used for data processing, statistical analyses, and figure generation are available on [GitHub](https://github.com/MandelliChristian) and will be made publicly accessible upon publication of the peer-reviewed version of this manuscript.

## Results

### Sequencing Quality Assessment for RNA-Seq and WGBS Data

RNA sequencing and whole-genome bisulfite sequencing (WGBS) datasets from both grapevine haplotypes demonstrated consistently high technical quality, ensuring reliable downstream analyses. For RNA-seq, all libraries exhibited excellent quality metrics, with average per-base Phred scores exceeding Q30 for over 95% of bases and total read counts ranging from approximately 31 to 41 million reads per sample. After trimming and quality filtering, more than 98% of reads were retained, GC content distributions centered around 44%, and 91.3–94.7% of reads mapped to the reference genome, of which over 85% mapping uniquely, confirming strong sequence integrity and alignment performance. WGBS data showed similarly robust quality across replicates. In Haplotype 1, alignment rates averaged 60.7% (59.9–60.7%), yielding ∼41.2 million aligned reads per sample. After deduplication, 29.5 million unique reads were retained, yielding an average sequencing depth of 15X and a DNA conversion efficiency of 99.2%. Methylation levels were consistent across replicates, with mCpG at 59.1–59.7%, mCHG at 28.8–29.6%, and mCHH at 3.4–3.8%. Haplotype 2 showed comparable results, with an average alignment rate of 60.2% (58.8–60.6%), ∼43.3 million aligned reads per sample, 34.2 million unique reads after deduplication, an average sequencing depth of 16X, and a 99.3% conversion efficiency. Its methylation levels ranged from 56.9–58.1% for mCpG, 28.7–29.4% for mCHG, and 3.4–3.8% for mCHH. Low duplication rates (0–18%) and consistent read lengths of 101 bp across both haplotypes further confirmed the reliability, reproducibility, and suitability of these sequencing datasets for integrated transcriptomic and epigenomic analyses.

### Haplotype-Specific Transcriptional Dynamics Reveal Divergent Strategies of Viral Response

After performing the quality control of the RNA-Sequencing, we processed the data to identify Differentially Expressed Genes (DEG) by comparing the transcriptome of infected versus control plants. Control plants were Agrobacterium-infected plants without the virus. We identified in both haplotypes around 9,000 total DEG on average, during the first two collection time points, but this total number significantly dropped to ∼2,000 genes at the third collection time (12 dpi) (Fig 1a), probably indicative of a transcriptional readjustment toward native or recovery conditions following the initial stages of vital infection. For unique DEGs, the relative proportions of up and down-regulated genes were also equally distributed across the three time points (Fig. 1b). Using a high-level confidence allele-pair dataset across the two haplotypes, we found that up to ∼77% of total DEG had paired alleles; a negligible number of genes was found to have an opposite regulation (up versus down), when compared to the corresponding allele in the other haplotype (Fig.1c). The proportion of unpaired alleles was ranging from 23 to 28% depending on the stage, which could overall explain the Gene Ontology (GO) enrichment differences observed between haplotypes (Figure S1). Using the up and down category of DEG per time point, we found that a GO enrichment in Haplotype 1 (H1) exhibited RNA metabolism, vesicle-mediated transport, and RNA stability pathways, with strong chloroplast-, plastid-, and mitochondria-related signatures along with an enrichment for RNA- and protein-binding, ubiquitin transferase, and GTPase activities (Fig. S1a). By contrast, Haplotype 2 (H2) showed enrichment for gene sets associated with cellular organization, nuclear and chromatin-associated complexes, and regulatory molecular functions, including binding and kinase activity (Fig. S1b), suggesting a response dominated by transcriptional and cell-cycle–related reprogramming rather than photosynthetic or plastid-associated processes. From this first gene set enrichment analysis, we observed evidence of a haplotype-specific response in the transcriptome following infection.

**Figure 1.**
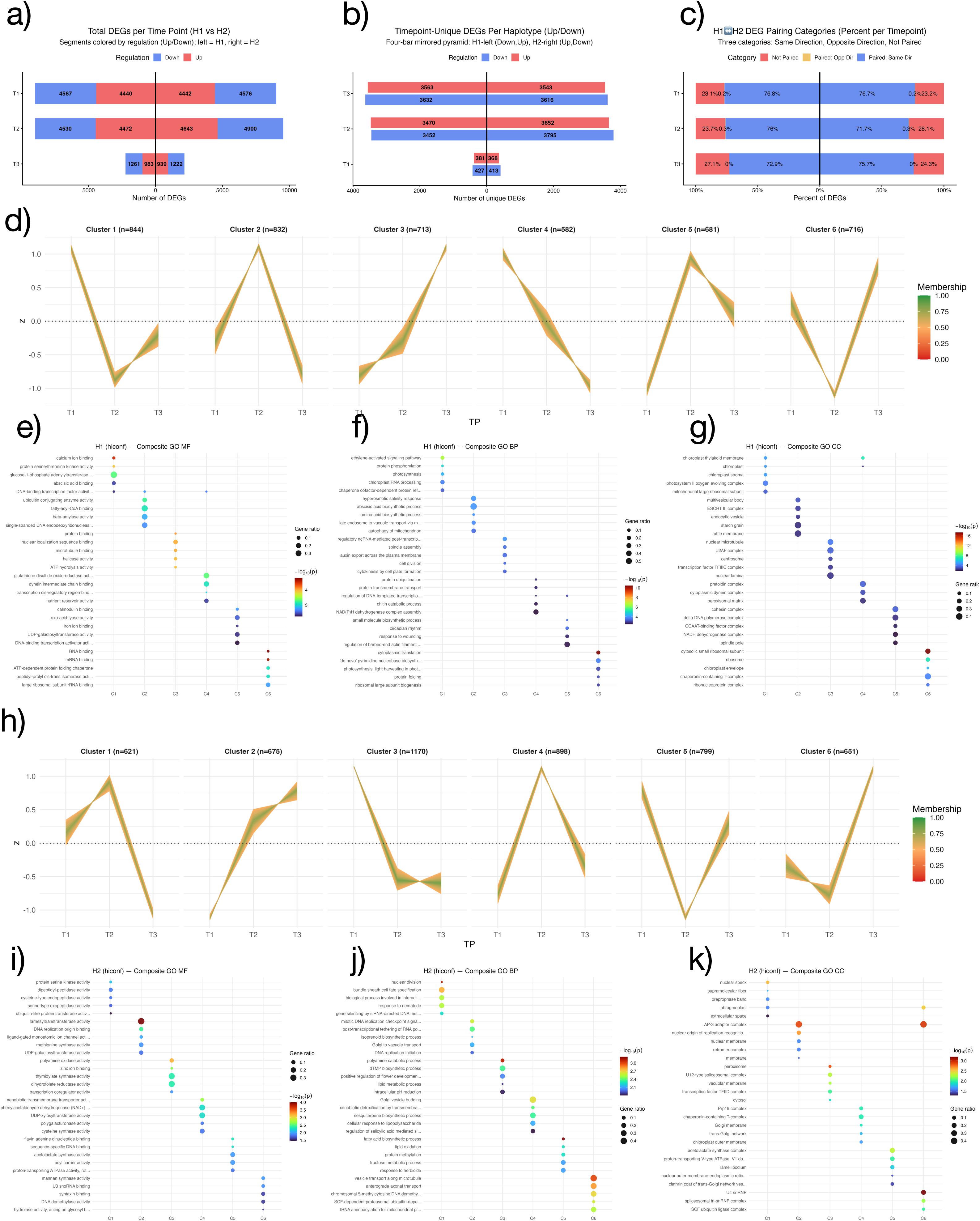
Overview of haplotype-specific temporal response patterns, functional enrichment, and soft-clustering structure. (a) Total differentially expressed genes (DEGs) per haplotype (H1, H2) across infection time points. (b) Temporal distribution of unique DEGs per haplotype (Up/Down regulated). (c) H1↔H2 pairing categories (paired, unpaired, and paired with opposite regulation) summarized per time point. (d–f) Composite GO enrichment for H1 clusters (BP, CC, MF), showing the top 5 functional signatures associated with each major temporal expression pattern. (g) Soft-clustering profiles for H1 (k = 6), displaying the membership-weighted mean trajectories across infection stages (T1–T3). (h–j) Composite GO enrichment for H2 clusters (BP, CC, MF). (k) Soft-clustering profiles for H2 (k = 6), capturing distinct temporal expression dynamics relative to H1.

We further investigated this aspect of haplotype-specific transcriptional responses of the microvine by including a clustering expression analysis over the course of the experiment to examine whether a haplotype-specific gene set enrichment could be revealed. Using a soft-clustering method, we found that ∼35% of DEGs in both haplotypes exhibited a pronounced transcriptional response and could be clustered into six major co-expression patterns per haplotype (Fig. 1d,h). Only two pairs of co-expression patterns (Clusters 2,6-H1 with Cluster 4,5 - H2) showed a certain degree of co-expression similarity between the two haplotypes. The other four co-expression patterns in H1 shared no expression similarity with those remaining in H2. A GO enrichment analysis, conducted in each cluster of the two haplotypes, revealed a distinct enrichment per co-expression pattern with little to no overlap across co-expression patterns within a haplotype (Fig 1e,f,g and 1i,j,k). Likewise, the two co-expression patterns per haplotype that shared similar expression trends between the two haplotypes did not show similar GO term enrichment.

In H1, the six clusters encompassed 4,368 genes. Cluster 1 (n = 844) showed rapid and early activation, whereas Cluster 2 (n = 832) and Cluster 5 (n = 681) displayed transient mid-infection peaks linked to protein folding, chaperone activity, and DNA replication. Cluster 3 (n = 713) and Cluster 6 (n = 716) exhibited progressive upregulation toward T3, strongly enriched for translation, chloroplast organization, and photosynthesis-related processes.

Conversely, Cluster 4 (n = 582) was characterized by early downregulation of chromatin remodeling, transcriptional regulation, and mitotic genes, consistent with suppression of proliferation and nuclear activity during initial infection (Lukhovitskaya 2014, Petermann, 2018). Together, these dynamics suggest that H1 undergoes an early inhibition of cell-cycle processes, a mid-phase stress adaptation, and a late reactivation of translational and energy machinery. (Fig. S3).

In H2, the six clusters (Fig. S2b) included 4,814 genes. Clusters 1 (n = 621) and 4 (n = 898) exhibited temporary mid-infection activation, enriched for mitotic regulation, microtubule organization, and defense signaling. Cluster 2 (n = 675) and Cluster 6 (n = 651) showed consistent/late increases in biosynthetic and transport processes, including fatty acid, nucleotide, and amino acid metabolism, indicating a shift in metabolism during the later stages of infection. Conversely, Cluster 3 (n = 1,170) and Cluster 5 (n = 799) were strongly downregulated during mid-to-late infection, with significant enrichment for ribosomal, spliceosomal, and thylakoid components, suggesting suppression of translation and organellar energy production. Remarkably, these mid-to-late downregulated clusters also contained genes related to RNA silencing, siRNA-directed DNA methylation, and chloroplast signaling, suggesting that viral infection may result in the downregulation of post-transcriptional defenses and plastid-mediated immune responses (Fig. S3).

To further examine the haplotype-specific co-expression patterns among paired alleles, we rerun the clustering analysis by tracking down paired genes (the two alleles were defined with high confidence level) and unpaired genes (one allele per haplotype). We first optimized the clustering analysis to capture the maximum number of paired alleles (Fig. 2a). A membership of 0.5, along with ten identified co-expression patterns, was found to be the best compromise to maintain a relatively high number of paired alleles falling in the co-expression pattern (∼42% H1 -> H2, ∼41% H2 -> H1) with a minimal drift in the cluster centroid shapes (Fig S2). In addition, the majority of genes identified with a 0.75 cutoff remain in the same cluster as with a 0.5 cutoff, indicating that it did not introduce noise. In addition, each cluster pattern created in H1 has a counterpart in H2, suggesting the identification of high-confidence co-expression patterns within a core gene set with paired alleles.

**Figure 2.**
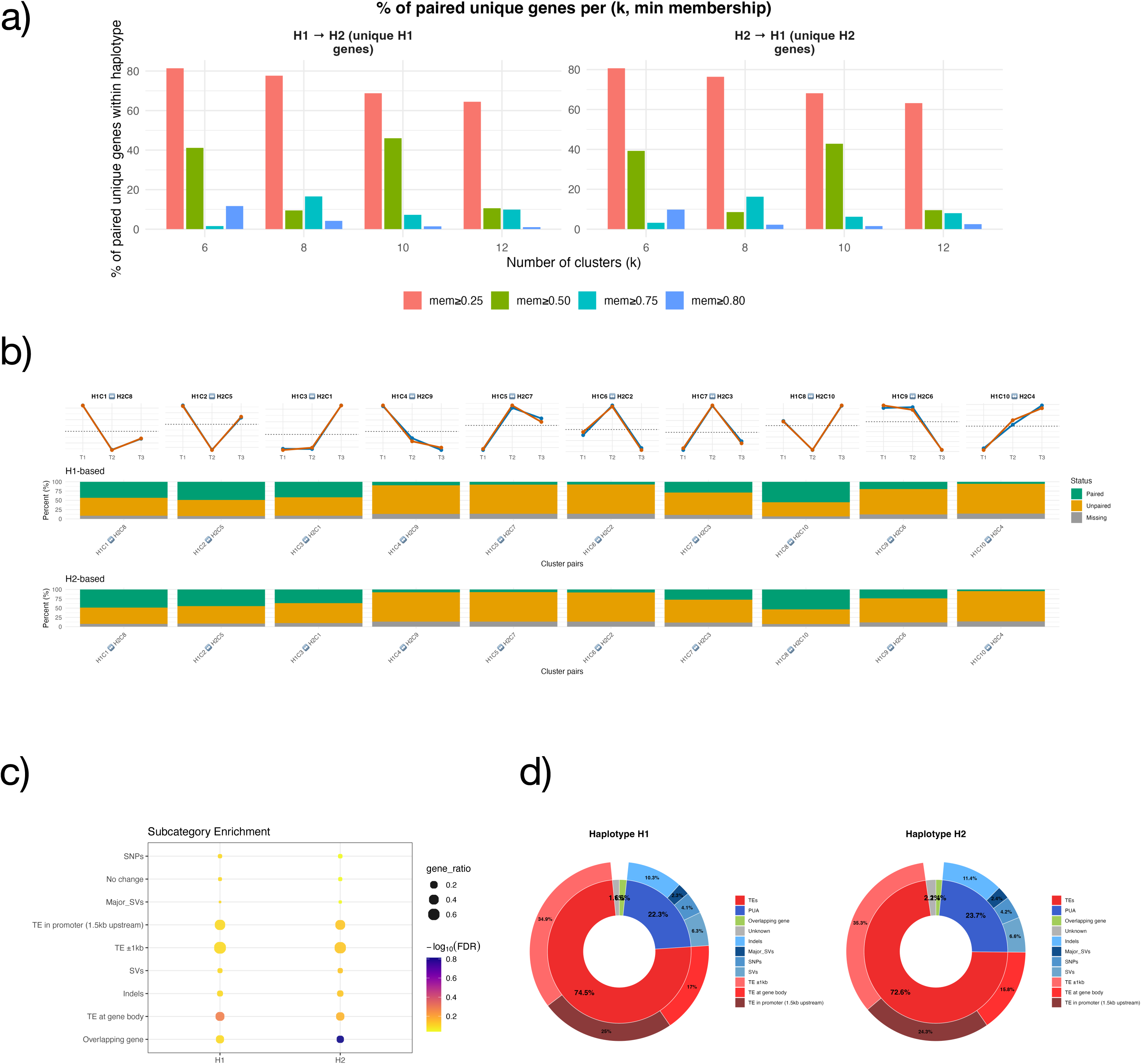
Multi-step analysis of haplotype-specific temporal clustering, pairing asymmetry, and functional divergence. (a) Proportion of unique DE genes in each haplotype that could be paired to a homologous transcript in the other haplotype across clustering resolutions (k = 6–12) and minimum membership thresholds (mem ≥ 0.25–0.80). Bars quantify the success of H1→H2 and H2→H1 pairings, illustrating haplotype-specific differences in cluster-level transcript correspondence. (b) Haplotype-resolved best-match cluster comparisons, showing temporal expression trajectories (top) and the proportion of paired versus unpaired genes for each cluster pair (bottom). (c) Enrichment of unpaired genes for structural, TE-associated, and regulatory categories, illustrating the genomic features most strongly associated with unpaired transcripts, lacking homologous temporal behavior across haplotypes. (d) Structural composition of H1 and H2 allele-specific expressed genes, represented as nested donut charts summarizing the contribution of major gene categories for each haplotype. (e) Composite GO enrichment for paired clusters (H1-C2↔H2-C1, H1-C5↔H2-C3, H1-C7↔H2-C2).

To ensure that this parameter choice is not influenced by replicate-level noise or over-partitioning, we further evaluated clustering robustness and structure. First, we assessed bootstrap stability by repeatedly resampling infected biological replicates within each time point (T1–T3) and rerunning the clustering. This analysis showed that cluster assignments at k = 10 and membership ≥ 0.5 are reproducible across bootstrap resampling (n = 50) of biological replicates, with a comparable proportion of genes retaining the same dominant cluster as observed for more stringent membership cutoffs (≥ 0.75 and ≥ 0.8). This is reflected by similar stability values (Fig. S4a and S4b) in the bootstrap plots at k = 10. Because stricter cutoffs (membership ≥ 0.75 and ≥ 0.8) did not improve reproducibility at this resolution but substantially reduced gene coverage, and because increasing resolution beyond k = 10 (k = 12) yielded no additional gain in stability and is associated with over-fragmentation, this combination (k = 10, membership ≥ 0.5) represents the highest resolution and least restrictive threshold that preserve robust cluster assignments. Second, we quantified centroid separation, defined as the mean pairwise distance between cluster centroids, to assess whether increased pairing occurred at the expense of distorted temporal expression patterns. As expected, centroid separation decreased smoothly with increasing k, reflecting finer subdivision of temporal programs; importantly, for any given k, centroid separation was highly consistent across membership thresholds (Fig. S4c and S4d). Lowering the membership cutoff to ≥ 0.5 did not induce an abrupt reduction in centroid separation relative to stricter thresholds, indicating that additional genes are incorporated without altering the underlying temporal trends. At higher resolution (k = 12), centroid separation continued to decline without revealing new distinct temporal structures, consistent with redundant subdivision rather than the discovery of biologically meaningful patterns. Together, these analyses independently support k = 10 and membership ≥ 0.5 as a robust and interpretable resolution for haplotype-aware temporal co-expression analysis.

Altogether, this set of co-expressed genes reflects a specific transcriptional response that exhibits a certain degree of haplotype specificity.

### Allele-centric co-expression analyses confirmed a Haplotype-Specific Transcriptome Remodeling During Infection

Following soft clustering with Mfuzz, six biologically enriched co-expression clusters per haplotype were identified (Fig. S2) and used as the foundation template dataset to assess temporal co-expression patterns among genes that shared expression in both alleles (paired alleles). To evaluate how consistently these behaviors were conserved across haplotypes, we quantified the percentage of paired genes assigned to the same cluster at varying clustering resolutions (k = 6–12) and membership thresholds (mem ≥ 0.25–1.00). The resulting bar plot (Fig. 2a) shows that the proportion of paired genes declines steeply as clustering resolution increases. At coarse granularity (e.g., mem ≥ 0.25), 60–80% of genes were shared between H1 and H2, indicating broadly similar infection-stage trajectories. However, at higher resolution (k = 10, mem ≥ 0.50), the overlap dropped below 50%, indicating that fine-scale temporal partitioning reveals extensive haplotype-specific co-expression.

Only a subset of cluster pairs (H1-C2 ↔ H2-C5, H1-C8 ↔ H2-C10) showed relatively strong reciprocal overlap (≥50%) (Fig. 2b), representing conserved temporal co-expression patterns that remain synchronized between H1 and H2. In contrast, the remaining cluster pairs (H1-C1↔H2-C8, H1-C3↔H2-C1, H1-C4↔H2-C9, H1-C5↔H2-C7, H1-C6↔H2-C2, H1-C7↔H2-C6, H1-C9↔H2-C5, H1-C10↔H2-C4) showed low haplotype pairing, highlighting extensive haplotype-specific expression.

Moreover, while most paired clusters exhibit tightly matched trajectories between H1 and H2, a subset of pairs (H1-C4↔H2-C9, H1-C5↔H2-C7, H1-C9↔H2-C5, H1-C10↔H2-C4) show markedly weaker overlap, consistent with haplotype-specific divergence in temporal regulation. Together, these trajectories underscore that although several infection-responsive programs are conserved, a subset is clearly haplotype-specific, potentially reflecting differences in stress adaptation and antiviral signaling efficiency between H1 and H2.

### TE and Promoter Variation Drive ASE and Functional Divergence of Antiviral and Stress-Response Modules Between Haplotypes

Before exploring the functional differences of the low-pairing clusters, we first assessed whether allele-specific expression in these genes might be connected to structural differences between alleles. Figure 2c and 2d show that unpaired ASE genes have strong, haplotype-specific enrichment for cis-regulatory changes. In H1, ASE mainly links to TE insertions within gene bodies, while in H2, ASE genes are enriched for promoter-proximal polymorphisms, including indels, SNPs, and major structural variants (SVs). These complementary enrichment patterns suggest that each haplotype has a distinct set of genomic features, likely predisposing different gene subsets to divergent expression. This structural asymmetry provides a mechanistic basis for the divergence in co-expression observed at the cluster level.

To gain functional insight into the eight cluster pairs that showed poor (<50%) reciprocal pairing between haplotypes (H1-C1↔H2-C8, H1-C3↔H2-C1, H1-C4↔H2-C9, H1-C5↔H2-C7, H1-C6↔H2-C2, H1-C7↔H2-C6, H1-C9↔H2-C5, H1-C10↔H2-C4), we performed GO enrichment separately for each cluster and summarized the results in composite dot plots (Fig. S3). Across all panels, clusters with similar temporal expression trajectories consistently resolved into haplotype-specific functional modules, indicating that low pairing reflects functional divergence rather than stochastic gene reassignment.

For the H1-C1 ↔ H2-C8 pair (Fig. S3a), the two haplotypes display a clear functional divergence despite broadly comparable temporal expression patterns. H1-C1 is predominantly enriched for chloroplast- and plastid-associated biological processes, including chloroplast RNA processing, plastid translation, photosynthesis-related functions, and multiple metabolic pathways linked to primary and secondary metabolism. These enrichments indicate a module focused on organelle-level metabolic regulation and plastid function, consistent with a coordinated adjustment of photosynthetic and biosynthetic activities during infection. In contrast, H2-C8 shows enrichment for a distinct set of functions emphasizing signaling, regulation, and protein-level control. Biological Process terms highlight hormone- and signal-mediated responses, defense-related pathways, and the regulation of protein phosphorylation and apoptosis. At the Cellular Component and Molecular Function levels, H2-C8 is further characterized by membrane-associated complexes, components of intracellular trafficking, and diverse regulatory activities, including enzyme modulation and protein–protein interactions.

For the H1-C3 ↔ H2-C1 pair (Fig. S3b), both clusters capture infection-responsive programs but diverge markedly in their regulatory emphasis. H1-C3 is enriched for processes linked to transcriptional and chromatin-level regulation, including chromatin organization, nucleosome assembly, epigenetic regulation of gene expression, and negative regulation of transcription. In contrast, H2-C1 exhibits a functional profile centered on protein turnover, stress-associated metabolism, and intracellular trafficking. Enriched categories include proteasome-related complexes, RNA polymerase and transcription-associated assemblies, membrane and vesicle-associated components, and enzymatic activities linked to carbohydrate and energy metabolism. This pattern indicates a shift toward post-transcriptional regulation and proteostasis, potentially reflecting enhanced protein quality control and stress buffering in H2. Thus, within this low-pairing module, H1-C3 prioritizes chromatin-based transcriptional regulation, whereas H2-C1 emphasizes protein-level regulation and metabolic adaptation.

For the H1-C4 ↔ H2-C9 pair (Fig. S3c), the two haplotypes separate along a pronounced structural–metabolic versus catabolic–stress response. H1-C4 is enriched for processes associated with cell wall organization, biosynthetic metabolism, and regulated phosphorylation, including cellulose microfibril organization, plant-type cell wall biosynthesis, and protein phosphorylation. These enrichments indicate a module focused on remodeling cellular architecture and coordinated biosynthetic activity, consistent with the reinforcement and restructuring of cell surfaces during infection. In contrast, H2-C9 shows enrichment for functions related to catabolic metabolism, organelle-associated respiration, and stress adaptation, with Biological Process terms emphasizing amino acid and lipid catabolism and nucleotide salvage, Cellular Component categories highlighting mitochondrial and plastid complexes, and Molecular Function enrichments dominated by oxidoreductase and dehydrogenase activities.

For the H1-C5 ↔ H2-C7 pair (Fig. S3d), the two haplotypes diverge in how they coordinate developmental regulation, transcriptional control, and cellular detoxification during infection. H1-C5 is enriched for processes linked to xenobiotic detoxification, transmembrane transport, transcriptional regulation, and growth- and development-related pathways, including circadian rhythm regulation, reproductive development, and peptide phosphorylation. At the Cellular Component level, H1-C5 emphasizes nuclear and transcription-associated complexes, while Molecular Function terms highlight DNA-binding transcription factor activity, histone reader functions, and RNA-processing enzymes, indicating a module centered on regulatory control and adaptive transcriptional reprogramming. In contrast, H2-C7 shows strong enrichment for vesicle trafficking, membrane-associated complexes, and intracellular transport, with prominent representation of clathrin-coated vesicles, plastid- and vacuole-associated compartments, and multiple transporter and hydrolase activities. These enrichments point to a functional emphasis on protein sorting, endomembrane dynamics, and cellular cleanup mechanisms.

For the H1-C6 ↔ H2-C2 pair (Fig. S3e), the two haplotypes diverge strongly in their engagement of autophagy- and nuclear stress–associated pathways. H1-C6 is enriched for processes related to organelle turnover and intracellular recycling, including autophagy of mitochondria, phagophore assembly, endosomal transport to the vacuole, and lipid and nitrate transport. These enrichments, together with terms linked to cellular respiration and responses to abiotic stress, indicate a module centered on metabolic reconfiguration and selective organelle degradation, consistent with active remodeling of cellular resources during infection. In contrast, H2-C2 shows enrichment for nuclear and protein-quality–control–associated functions, including chromatin- and transcription-related complexes, proteasome components, and multiple enzyme activities involved in ubiquitin-mediated modification and redox regulation. At the Cellular Component level, H2-C2 emphasizes nuclear and ribonucleoprotein assemblies, while Molecular Function terms highlight catalytic activities linked to protein processing and stress adaptation, suggesting a shift toward nuclear regulation and proteostasis rather than bulk autophagic turnover.

For the H1-C7 ↔ H2-C3 pair (Fig. S3f), the two haplotypes diverge in how they balance signaling, transport, and nuclear regulation during infection. H1-C7 is enriched for processes linked to membrane transport and hormone-associated signaling, including auxin-activated signaling pathways, fatty acid and amino acid transport, inositol metabolism, and plasmodesmata-mediated intercellular transport. These enrichments indicate a module centered on signal transduction and controlled molecular exchange, consistent with coordinated intercellular and hormonal responses. In contrast, H2-C3 is dominated by chromatin- and transcription-associated functions, including chromatin assembly, circadian regulation of gene expression, DNA replication–dependent chromatin organization, and multiple nuclear protein complexes. At the Molecular Function level, H2-C3 further emphasizes regulatory and enzymatic activities associated with transcriptional control and metabolic adjustment, indicating a shift toward nuclear reprogramming rather than membrane-level signaling.

The H1-C9 ↔ H2-C6 comparison (Fig. S3g) reveals divergence between signaling-driven stress responses and intracellular trafficking–centered regulation. H1-C9 is enriched for signaling pathways and regulatory processes, including cytokinesis-related regulation, kinase cascades, protein phosphorylation, and translational control, as well as metabolic processes such as galactose and lipid biosynthesis. This profile suggests a module that integrates signal transduction with metabolic and translational regulation. By contrast, H2-C6 shows strong enrichment for vesicle trafficking, membrane remodeling, and protein sorting, including multivesicular bodies, endomembrane compartments, and multiple transporter and enzyme activities. Together, this pair highlights a clear separation between regulatory signaling (H1) and cellular trafficking and turnover (H2) as alternative strategies for managing infection-associated stress.

Finally, for the H1-C10 ↔ H2-C4 pair (Fig. S3h), divergence emerges between regulatory and developmental control versus metabolic and enzymatic specialization. H1-C10 is enriched for processes linked to DNA ligation, RNA-mediated regulation, ribosome-associated quality control, and light-responsive and developmental transitions, indicating a module focused on regulatory coordination and transcriptional control during later stages of infection. In contrast, H2-C4 displays enrichment for metabolic enzyme activities and biosynthetic pathways, including carbohydrate metabolism, redox-associated enzymatic functions, and hormone-related biosynthesis. Cellular Component terms further emphasize the diversity of intracellular complexes, suggesting a shift toward metabolic execution rather than regulatory oversight.

Together, these eight comparisons demonstrate that genes sharing similar temporal expression trajectories are repeatedly partitioned into functionally distinct haplotype-specific modules. While a limited core of stress- and defense-related functions is shared, each haplotype consistently enriches additional, non-overlapping biological programs. These results indicate that fine-scale co-expression clustering resolves divergence in how H1 and H2 coordinate RNA regulation, chromatin dynamics, metabolism, and proteostasis during infection, revealing complementary yet mechanistically distinct strategies for coping with viral stress (Fig. S3a–h).

### WGCNA Reveals Haplotype-Specific Modules Associated with Viral Response and Chloroplast Function

WGCNA identified several modules of co-expressed genes whose eigengene dynamics and internal connectivity patterns were markedly influenced by GRBV infection. Comparative analysis between haplotypes revealed paired modules with substantial gene overlap (>60%) and conserved eigengene trends, yet also distinct infection-dependent rewiring of network relationships.

Among these, the blue module in haplotype 1 (H1-Blue) and its paired purple module in haplotype 2 (H2-Purple) (Fig. S5b) were particularly notable. These modules were enriched in chloroplast- and plastid-associated genes with hydrolase activity (Fig. 3c – left panel) and in genes related to cell differentiation and developmental regulation (Fig. 3c – right panel), respectively, suggesting that they may participate in chloroplast-linked developmental and regulatory processes modulated during infection.

**Figure 3.**
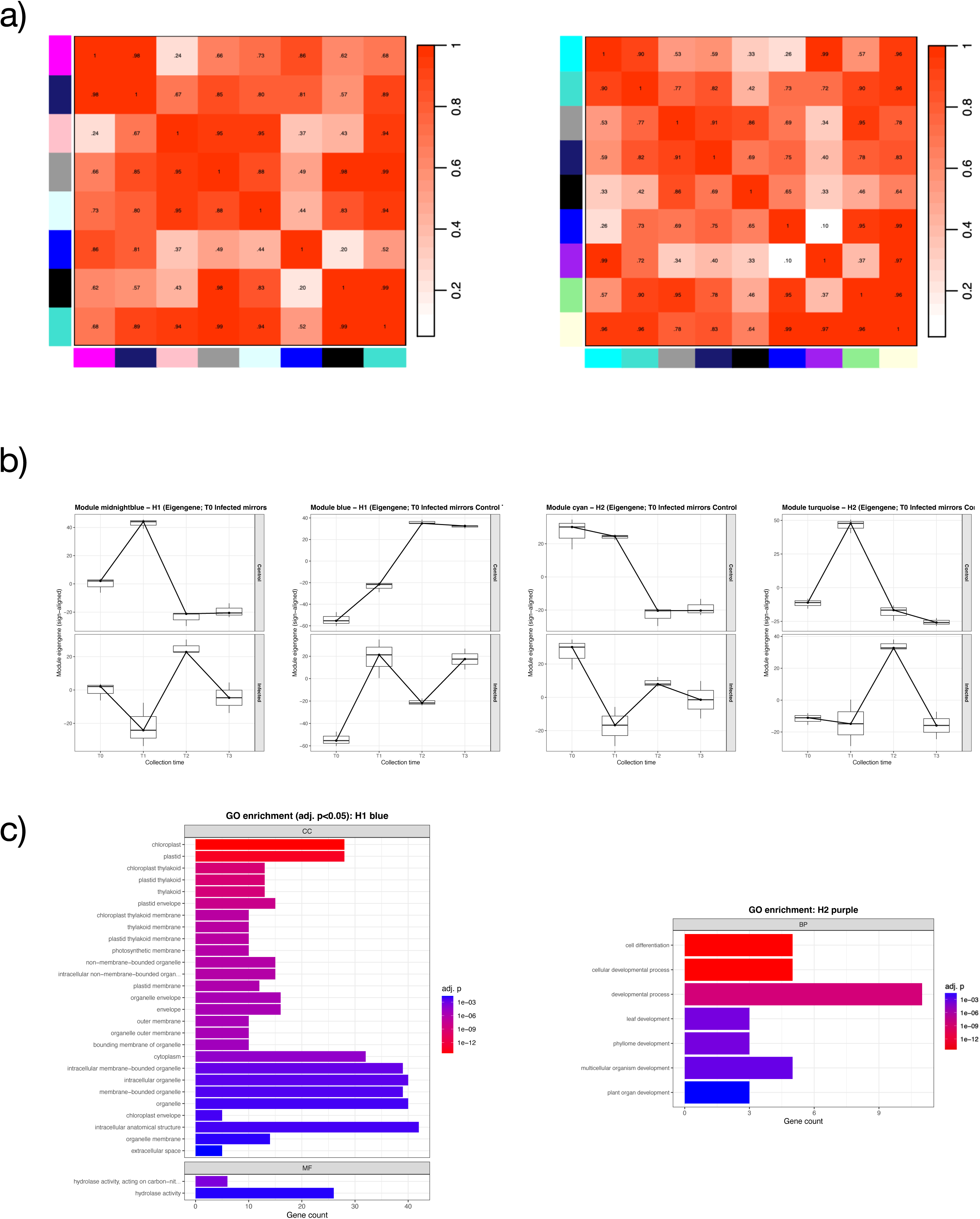
WGCNA modules preservation and functional characterization of key modules. (a) Modules preservation heatmaps for haplotypes H1 (left) and H2 (right), showing the strength and significance of preservation (bicor) between modules in infected and control samples. Warmer colors indicate stronger positive correlations, whereas cooler colors represent negative or weak preservation. (b) Eigengene expression profiles of the most divergent modules in each haplotype, illustrating characteristic temporal trajectories and highlighting modules with early, mid, or late activation or repression during infection. (c) Functional enrichment (GO Biological Process) for paired modules exhibiting strong temporal dynamics or haplotype-specific behavior (e.g., H1–blue, H2–purple). Barplots display significantly enriched terms (adj. p ≤ 0.05) and the number of genes associated with each function.

In H1-Blue, the hub gene Vitvimv05.H1_g10577 (peptidyl-prolyl cis–trans isomerase; kME = 0.984) emerged as a strong candidate for viral interaction. A homolog in *Nicotiana benthamiana* has been identified as an interactor of the Tomato leaf curl New Delhi virus movement protein (Chang et al. 2022), which suppresses the isomerase’s defense functions through direct physical interaction and transcriptional inhibition. This mechanism facilitates viral movement and infection, suggesting that Vitvimv05.H1_g10577 could play a similar role in GRBV susceptibility.

Interestingly, a homologous gene on haplotype 2 (Vitvimv14.H2_g28970, peptidyl isomerase; kME = 0.926) also functions as a hub gene in the H2-Purple module, indicating conserved roles across haplotypes. In both modules, eigengene (ME) vectors showed a noticeable decrease at T2 in infected samples compared to controls, reflecting a temporary suppression of chloroplast-related transcription during mid-infection. Another hub gene in H2-Purple, Vitvimv12.H2_g22850 (NADPH–protochlorophyllide oxidoreductase), strengthens the connection to chloroplast function and energy metabolism. Genes associated with chloroplasts are known to influence viral genome replication and host defense regulation, as viral replication complexes often anchor to host chloroplast membranes (Shahriari et al. 2025). The enrichment of hydrolase activity within these modules further supports their role in viral DNA binding and pathogenesis (Gnanasekaran et al. 2023).

Furthermore, no hub genes were shared between H1-Blue and H2-Purple despite their strong gene-level overlap, indicating that each haplotype may depend on different central regulators within otherwise similar co-expression networks. Notably, in all paired modules (>60% shared genes), the hub genes were non-allele-specific (non-ASE), suggesting that expression divergence across haplotypes occurs at the modular level rather than through ASE effects on individual hub genes.

Another set of paired modules, H1-Black and H2-Blue (Fig. S5a), showed high structural similarity and shared two hub genes (Fig. S6a). Among them, Vitvimv06.H1_g12518 (calcium-binding protein) is particularly notable. In *Nicotiana*, calcium-binding proteins have been shown to interact with geminivirus V2 proteins, and calcium signaling has been linked to RNAi-based antiviral defense (Wang et al. 2021). This suggests a potential regulatory role for Vitvimv06.H1_g12518 in calcium-mediated antiviral responses to GRBV. Other shared hub genes, including Vitvimv12.H1_g24876 | Vitvimv12.H2_g24032 (ATPase) and Vitvimv13.H1_g25175 | Vitvimv13.H2_g24288 (nematode-resistance protein), are functionally related to host and viral DNA replication and the positive regulation of viral resistance, respectively. The intra-modular relationships among these shared hub genes seem to be conserved but relatively weak, with connections represented by thin, distant edges in the network (Fig. S6a). This may indicate functional conservation with some divergence in co-expression strength.

Infection significantly altered inter-modular relationships, causing a widespread loss of eigengene correlations within haplotypes (Fig. 3a). In haplotype 1, H1-Blue and H1-Black, which were positively correlated under control conditions, lost their connection after infection, showing distinctly different ME-vector trajectories (Fig. 3b – right panels). Likewise, H1-Blue experienced a loss of connectivity with the H1-Pink and H1-Lightcyan modules. This disruption likely indicates a breakdown in coordinated transcriptional programs governing chloroplast activity and regulation.

The H1-Pink module also displayed reduced preservation, losing significant relationships with H1-Magenta and H1-Black during infection, suggesting broad-scale rewiring of gene networks in response to viral stress.

A comparable pattern was observed in haplotype 2. The paired H2-Purple module, which mirrors H1-Blue, lost its relationships with H2-Grey60, -Blue, -Black, -Midnightblue, and - Lightgreen modules. This pattern of inter-modular loss of relationship was consistent across haplotypes, implying that infection triggers a coordinated yet haplotype-independent loss of network preservation in chloroplast-related modules.

The simultaneous loss of inter-modular connectivity in H1-Blue and H2-Purple underscores that these modules, which share genes and hub-gene functions, are among the most infection-sensitive areas of the transcriptome. Although they show similar eigengene trajectories (generally increasing over infection timepoints), both modules diverged from the rest of the co-expression network under viral pressure. These results indicate that GRBV infection specifically disrupts chloroplast-related defense hubs that are otherwise conserved across haplotypes, possibly reflecting an adaptive viral strategy to target plastid pathways and weaken host defenses (Bhattacharyya et al. 2015; Zhao et al. 2023, 2024).

### Transposable Element–Linked and Promoter Methylation Drive Haplotype-Specific Epigenetic Divergence During GRBV Infection

WGBS was performed on both haplotypes to generate single-base resolution DNA methylation maps. Global methylation levels were highly similar between haplotypes across all cytosine contexts (CG, CHG, CHH), indicating that the overall methylome architecture is largely conserved (Fig. 4a).

**Figure 4.**
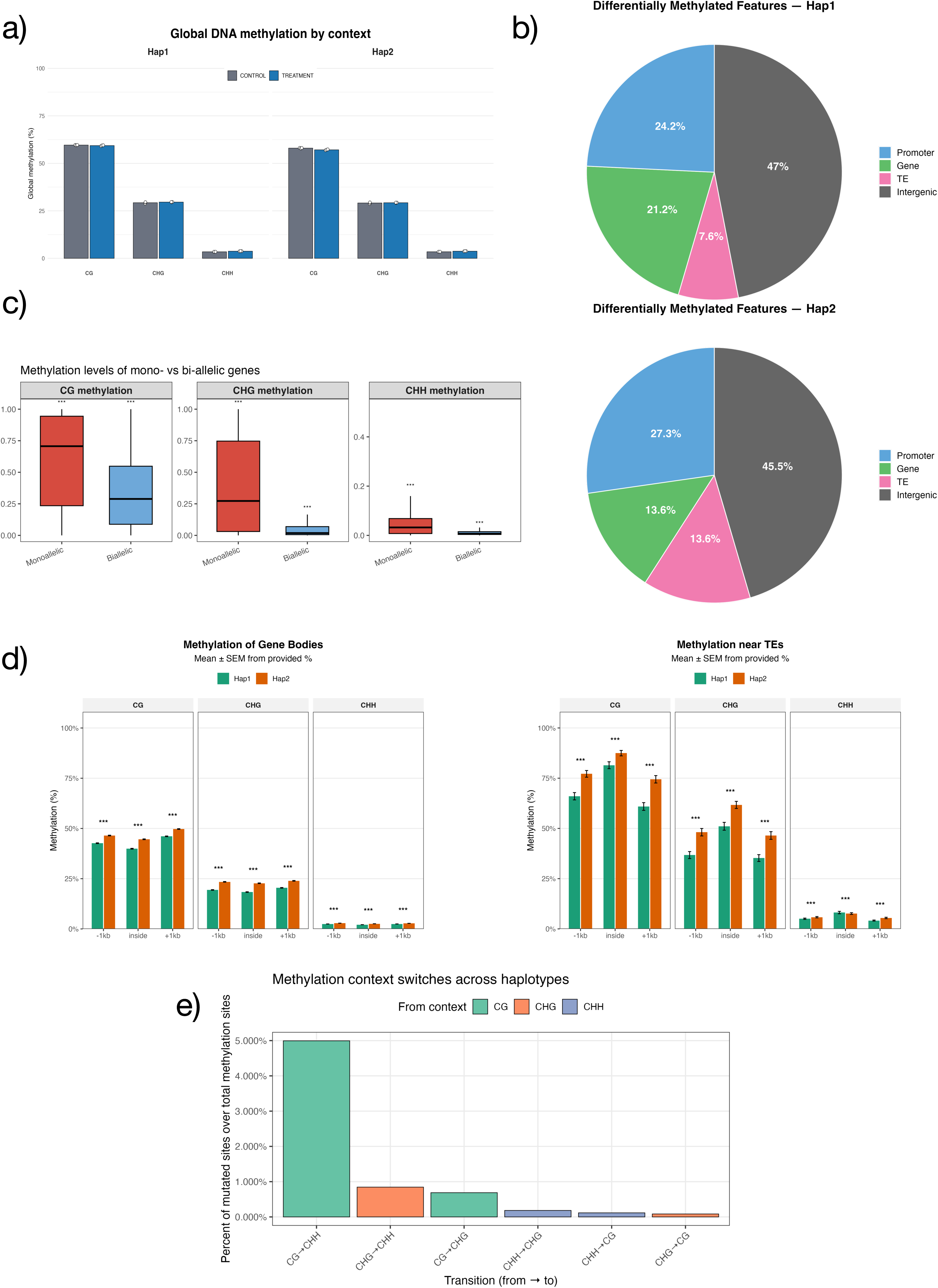
Haplotype-resolved DNA methylome and GRBV-induced epigenetic reprogramming in Microvine V4. (a) Whole-genome methylation levels of haplotype 1 (Hap1) and haplotype 2 (Hap2). (b) Ratio of Differentially Methylated Features (DMFs) of Hap1 (upper panel) and Hap2 (lower panel). (c) Methylation levels of mono- and bi-allelic genes. (d) Methylation level of gene bodies (left), TEs (right), and of the upstream and downstream 1 kb regions. (e) Bar plot showing the relative frequency of cytosine-context transitions between haplotypes. Percentages are normalized by the total number of source-context sites (e.g., total CG sites in the reference haplotype) and aggregated across both haplotypes. The height of each bar represents the proportion of cytosines that changed methylation context.

The genomic distribution of differentially methylated features (DMFs) (Fig. 4b) highlighted distinct epigenetic architectures between haplotypes. In Haplotype 1 (H1) (Fig. 4b – upper panel), DMFs were enriched in intergenic regions (47%), with 7.6% overlapping intergenic TEs, 24.2% located in promoters, and only 21.2% within gene bodies, indicating that most methylation variation occurs in promoter or intergenic regions. By contrast, Haplotype 2 (H2) (Fig. 4b, lower panel) displayed a relative increase in promoter-associated DMFs and a corresponding reduction in intergenic fractions, together with nearly a two-fold enrichment of TE-associated DMFs, reflecting a shift toward both promoter and transposon-linked methylation. These distributions show that methylation levels of TE-linked and distal regulatory elements are the major sources of epigenetic divergence between the two haplotypes.

To explore the relationship between methylation divergence and allele-specificity, we examined methylation of mono- and bi-allelic genes across haplotypes (Fig. 4c). Monoallelic genes (genes with one allele present in only one of the two haplotypes) in our dataset exhibited significantly higher methylation within gene bodies than biallelic genes (genes with one allele for each haplotype). These results indicate that DNA methylation asymmetries between haplotypes are closely coupled to allelic expression imbalance, suggesting that epigenetic divergence contributes to the regulation of haplotype-specific transcription under GRBV infection.

Analysis of methylation profiles across genes (Fig. 4d, left panel) and TEs (Fig. 4d, right panel) revealed clear differences between haplotypes. In both H1 and H2, CG methylation was highest within gene bodies, while CHG methylation had similar methylation levels in flanking regions and TEs. However, compared with H1, H2 showed a global increase in methylation across all contexts.

To quantify cytosine-context reconfiguration directly, we aligned the two haplotypes and assessed all single-base cytosines that shifted context between H1 and H2 (Fig. 4e), as previously reported (Zhong et al. 2022a). Most transitions involved the conversions between CG -> CHG and CHG -> CHH contexts. Together, these results demonstrate that although global methylation levels are conserved, local cytosine-context transitions and TE-linked methylation highlight extensive haplotype-specific epigenetic divergence under GRBV infection.

Hierarchically clustered heatmaps of the top differentially methylated genes (DMGs) revealed highly consistent methylation reprogramming across biological replicates in response to GRBV infection, with infected samples clearly separating from controls in both haplotypes (Fig. S7). In Haplotype 1 (H1) (Fig. S7a), the strongest methylation shifts were observed on two unannotated loci (Vitvimv01.H1_g1417 and Vitvimv05.H1_g10598; both hypermethylated), two adjacent ATP-dependent RNA helicases (Vitvimv13.H1_g26075 and Vitvimv13.H1_g26079; both hypomethylated), and an oxidative-stress enzyme, NADH-cytochrome b5 reductase (Vitvimv06.H1_g12008; hypermethylated). These results suggest that GRBV infection in H1 primarily remodels RNA-metabolic and redox-associated pathways.

In contrast, Haplotype 2 (H2) (Fig. S7b), exhibited hypermethylation of an unannotated gene (Vitvimv01.H2_g1241), the developmental regulator EDA18 (Vitvimv01.H2_g1518), and glutathione S-transferase GSTU10 (Vitvimv01.H2_g1301), a key component of oxidative defense and plant–pathogen responses (Gullner et al. 2018). The strongest hypomethylation occurred at Vitvimv03.H2_g5456 (unannotated) and Vitvimv04.H2_g7667 (DHHC-type zinc-finger protein), indicating modulation of signaling and regulatory processes. Interestingly, the ABC transporter G family member 22 (Vitvimv13.H1_g26109 and Vitvimv13.H2_g25189) was the only gene exhibiting differential methylation across both alleles.

Notably, every differentially methylated gene (DMG) contained a transposable element (TE) either within its gene body or within 1 kb of its flanking regions. This pattern aligns with recent reports (Zhong et al. 2022b) showing that differentially methylated cytosines (DMCs) are preferentially located within TEs and distal intergenic regions, whereas they are relatively depleted in gene bodies and proximal flanks, suggesting that TEs represent hotspots for DNA methylation changes.

These findings indicate that although global methylation levels remain comparable across haplotypes, GRBV infection induces extensive, yet haplotype-specific, epigenetic reprogramming. In H1, the response is characterized by broad methylation changes affecting RNA metabolism and redox-associated pathways outside promoter regions. In contrast, H2 displays focused hypermethylation targeting oxidative-stress and developmental regulators, accompanied by a pronounced enrichment of promoter- and TE-associated methylation. These divergent methylation responses underscore distinct epigenetic strategies governing host–virus interactions across haplotypes.

## Discussion

By separating the two parental haplotypes, phased genomes reveal regulatory differences, allelic diversity, and stress-response pathways that would otherwise be hidden in collapsed assemblies. In *Vitis vinifera*, a species marked by high heterozygosity and cultivar-dependent disease responses, a haplotype-resolved genome is therefore crucial for uncovering how distinct alleles shape the plant-pathogen interactions.

Our integrative transcriptomic and temporal clustering analyses showed that the two haplotypes employ fundamentally different response strategies to GRBV infection. H1 showed early suppression of nuclear, cell-cycle, and chromatin-related processes, followed by a gradual reactivation of translational, metabolic, and photosynthetic pathways (Fig. 1d). This pattern suggests a tolerance-like approach, in which the host temporarily restricts growth and cellular activity during peak viral pressure but restores homeostasis once replication stabilizes. Conversely, H2 showed a less sustained defense response, characterized by early activation of stress, immune, and hormone-signaling pathways, along with significant viral suppression of RNA silencing and chloroplast-related signaling (Fig. 1e). The unsustained activation of defense pathways, coupled with impaired recovery of chloroplast defense modules, suggests that H2 experiences a more pronounced breakdown of antiviral coordination.

Both haplotypes exhibited a transition from an initially stochastic transcriptional phase, marked by heterogeneous and variable expression patterns, to a later stabilized phase with fewer DEGs (Fig. 1a). This early-to-late progression mirrors patterns observed in mammalian viral infections, in which intrinsic noise in signaling pathways drives early response variability before converging on stable steady states (Blake and Collins 2005; Zhao et al. 2012). In our system, this shift likely reflects the establishment of metabolic tolerance in H1 and chronic stress adaptation in H2.

Given these contrasting transcriptional responses, we next asked whether epigenetic regulation contributed to the divergence in haplotypes’ responses. Whole-genome methylation profiling confirmed that each haplotype exhibits a distinct regulatory landscape during infection, with differences concentrated in promoters and TE-rich regions.

H1 showed changes in methylation across distal regulatory and intergenic regions, whereas H2 displayed a more targeted hypermethylation, focused directly on promoters and TE-rich loci. These TE-linked methylation shifts likely arise from sequence-dependent differences in transcription factor (TF) and insulator binding (Wei et al. 2025), including variation at CTCF, KRAB-ZNF, and other cis-regulatory motifs known to modulate local chromatin states (Do et al. 2016). Because TEs are major hotspots of DNA methylation and epigenetic surveillance (Zhong et al. 2022a), their enrichment among differentially methylated features (Fig. 4b and 4d) suggests they may serve as epigenetic sensors that enhance haplotype-specific responses under viral stress. Overall, these patterns indicate that DNA methylation can either buffer or amplify regulatory signals during infection in a haplotype-dependent manner. In H1, the broader remodeling of distal regions may support long-term regulatory stability and adaptive immunity. This reflects an epigenetic strategy that leverages dynamic promoter methylation/demethylation, as observed in plants, where RNA-directed DNA methylation (RdDM) and active promoter demethylation coordinate gene expression during development, viral and environmental stress responses (Sato et al. 2017; Zhang et al. 2018).

In contrast, the promoter- and TE-focused hypermethylation seen in H2 appears to be a more reactive epigenetic defense mechanism that may ultimately impede RNA silencing and immune coordination. This is especially true, as TEs are usually silenced to protect genomic stability and reduce inflammation. Yet the host might deliberately derepress them during viral infection to enhance antiviral signaling (Hale 2022).

This interaction between TE-associated methylation and allele-specific regulation highlights the broader role of epigenetic flexibility in shaping different viral responses among heterozygous genomes. High-throughput methylome profiling (Arora et al. 2022) further supports this, revealing stress- and cultivar-specific DNA methylation patterns. Notably, in rice, single-base-resolution methylome analyses under abiotic stress have shown that cultivar-specific differences in gene expression are linked to distinct DNA methylation patterns, with either hypo- or hypermethylation associated with increased expression of stress-responsive genes. More importantly, these epigenetic differences often arise from underlying SNPs, indicating that haplotype-specific sequence variation can drive differential methylation and shape divergent regulatory responses (Rajkumar et al. 2020). Similarly, in lemon, allele-specific DNA methylation at the PH5 promoter explained the contrasting citric acid levels between cultivars (Wang et al. 2025). Together, these findings underscore how genetic variation and TEs dynamics converge through epigenetic mechanisms to enable fine-tuned, haplotype-specific responses to environmental and biotic challenges. This resolution over haplotype-specific methylation not only reveals key regulatory mechanisms but also paves the way for functional validation of allele-specific epigenetic marks, which may accelerate precision breeding and the development of cultivars with enhanced stress resilience and optimized traits.

Having established that transcriptomic and methylation dynamics sharply diverge between haplotypes, we then analyzed how these regulatory differences appear at the level of co-expression network organization, especially within biological processes known to be influenced by geminivirus infection.

Both haplotypes experienced pronounced GRBV-mediated disruption of chloroplast-centered co-expression modules identified through WGCNA. Modules such as H1-Blue and H2-Purple lost inter-modular connectivity and showed transient reductions in eigengene expression during mid-infection (Fig. S5b), indicating that chloroplast function constitutes a shared point of vulnerability across haplotypes. However, following soft clustering, the extent and outcome of this disruption varied significantly between the two haplotypes. In H1, chloroplast-related genes within Mfuzz clusters 3 and 6 (Fig. S2a) were upregulated during the later stages of infection, indicating a recovery phase marked by the restoration of photosynthetic and plastid metabolic functions. In contrast, H2 exhibited a sustained decline in chloroplast defense and immune coordination, including reduced expression of key hub genes such as peptidyl-prolyl isomerases and NADPH-protochlorophyllide oxidoreductase. Together, these patterns support a model in which GRBV replication complexes preferentially interfere with plastid-based defense signaling, compromising both energy metabolism and plastid-derived antiviral responses. The more persistent network deterioration observed in H2 likely contributes to its weaker ability to recover functional chloroplast activity following viral pressure.

Building on this network-level disruption, we next examined how temporal regulation of RNA silencing and DNA methylation pathways contributed to haplotype-specific infection trajectories. Temporal clustering showed that H1 reactivated core RNAi components during the later stages of infection, aligning with its recovery of chloroplast metabolic activity and supporting a coordinated reinstatement of antiviral defenses once the viral load stabilized. In contrast, H2 exhibited a mid-infection suppression of both RNAi- and methylation-related genes, a pattern that overlapped with peak viral replication and further disrupted chloroplast immune coordination. This aligned suppression of epigenetic and silencing pathways likely compromises H2’s ability to contain GRBV infection, reinforcing the transcriptomic and methylation evidence that it mounts a more reactive yet less sustained antiviral response.

Together, the integration of transcriptomic, “methylomic”, and network dynamics suggests that H1 and H2 follow different regulatory trajectories in responding to GRBV: H1 gradually transitions from early suppression to controlled metabolic recovery supported by late-stage reactivation of RNAi defenses, whereas H2 experiences a compounding disruption of methylation, RNA silencing, and plastid immunity that limits its capacity to regain homeostasis. These multi-layered patterns highlight how haplotype-resolved genomics can uncover divergent antiviral strategies that would be obscured in collapsed analyses, providing a deeper understanding of how allelic variation shapes disease outcomes in heterozygous perennial hosts.

## Abbreviations

ASE: allele-specific expression
bp: base pair
CDS: coding sequence
DEG: differentially expressed gene
DMF: differentially methylated feature
DMG: differentially methylated gene
dpi: days post infiltration
GRBV: Grapevine red blotch virus
GO: Gene Ontology
H1: haplotype 1
H2: haplotype 2
kME: module eigengene-based connectivity
ME: module eigengene
RNAi: RNA interference
RNA-seq: RNA sequencing
SV: structural variant
TE: transposable element
TOM: topological overlap matrix
vsiRNA: virus-derived small interfering RNA
WGBS: whole-genome bisulfite sequencing
WGCNA: weighted gene co-expression network analysis

## Acknowledgments

We thank Keith Lloyd Perry (Cornell University -) for providing the bitmer construct containing the GRBV genome. We also thank the California Department of Food and Agriculture (CDFA) and the Oregon Wine Research Institute (OWRI) for their financial contribution to the study.

