## Supplemental Figures for "Grapevine Red Blotch Virus Induces Haplotype-specific Genetic and Epigenetic Responses"

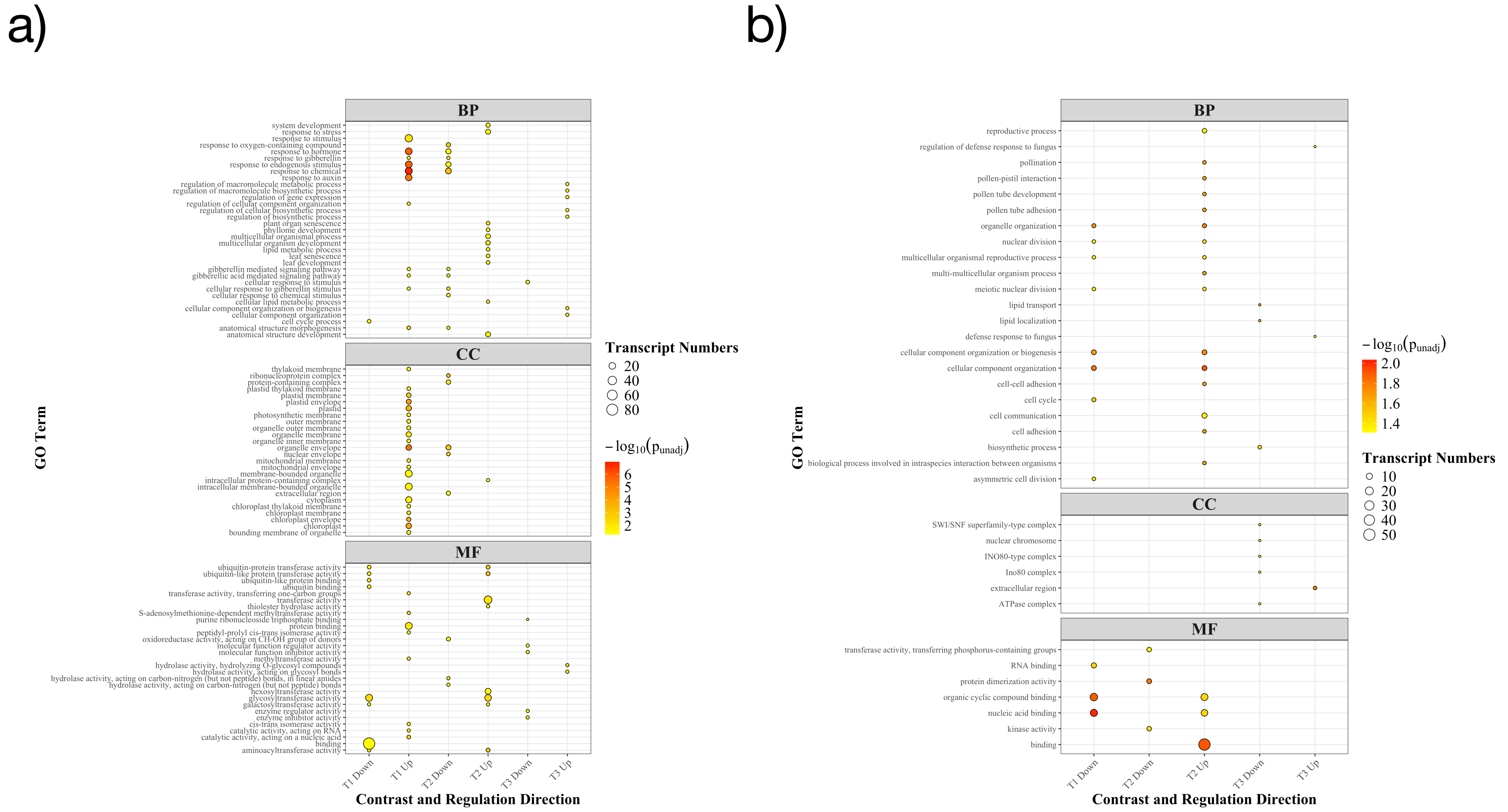


**Figure S1. Functional enrichment of differentially expressed transcripts across contrasts and regulatory directions.** GO enrichment results (BP, CC, MF) for each contrast and regulation direction (Up/Down), shown as bubble plots. Each point represents a significantly enriched GO term, with bubble size proportional to the number of DE transcripts associated with the term and color indicating statistical significance (–$\log_{10}$ p-value). Panels summarize how biological processes (BP), cellular components (CC), and molecular functions (MF) are differentially activated or repressed across infection stages. The left panel a) represents the enrichment plot for H1, the right panel b) for H2.

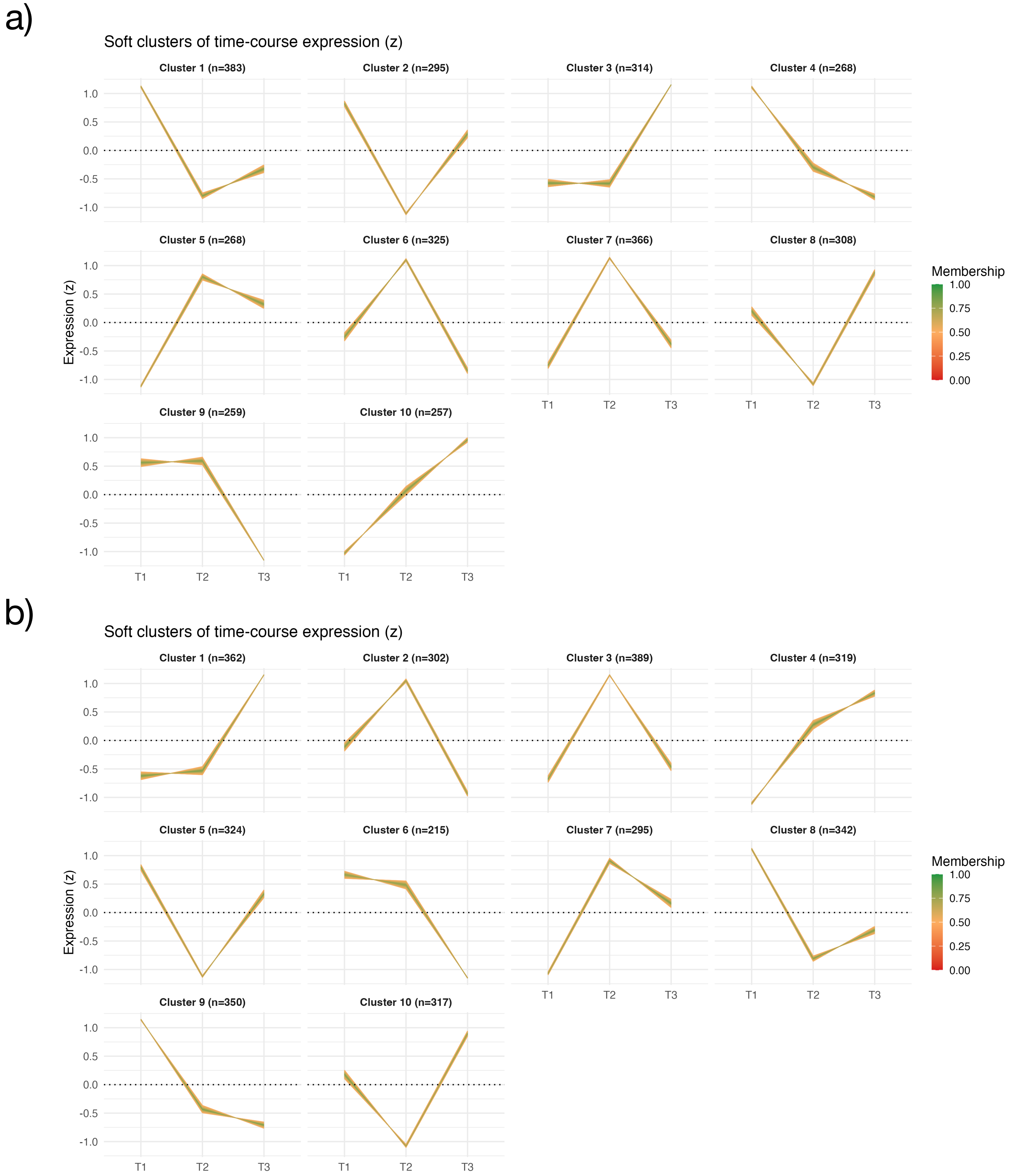


**Figure S2. Soft clustering of temporal expression profiles across infection.** (a) Haplotype H1: Mfuzz soft clustering (k = 10) of DE genes detected at any infection time point, illustrating major temporal expression trajectories across T1–T3. Each panel shows a cluster-specific mean trajectory (line) overlaid with all member gene profiles colored by membership strength (0–1), reflecting confidence in gene–cluster assignment. (b) Haplotype H2: Equivalent soft-clustering results (k = 10) revealing shared and haplotype-specific temporal patterns relative to H1.

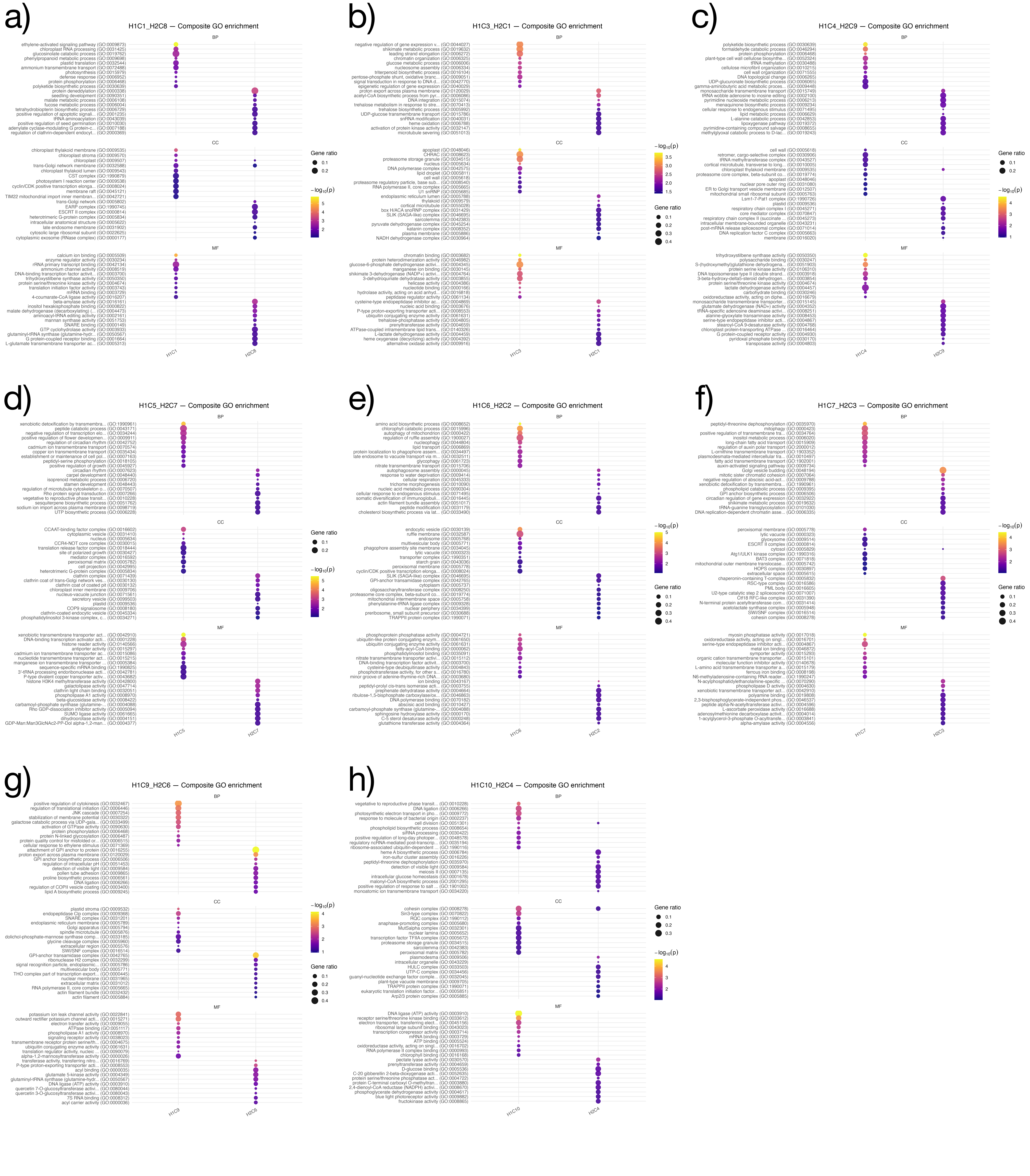


**Figure S3. Composite GO enrichment for paired clusters (H1-C1↔H2-C8, H1-C3↔H2-C1, H1-C4↔H2-C9, H1-C5↔H2-C7, H1-C6↔H2-C2, H1-C7↔H2-C6, H1-C9↔H2-C5, H1-C10↔H2-C4).** Bubble plots show Gene Ontology (GO) enrichment results obtained from composite gene sets formed by reciprocally paired Mfuzz clusters between haplotypes H1 and H2 that exhibit low inter-haplotype gene pairing. For each cluster pair, enriched GO terms are displayed across the three main ontologies, Biological Process (BP), Cellular Component (CC), and Molecular Function (MF). Dot size represents the number of genes (as a ratio) contributing to each GO term, while color denotes enrichment significance ($-\log_{10}$ adjusted *p*-value).

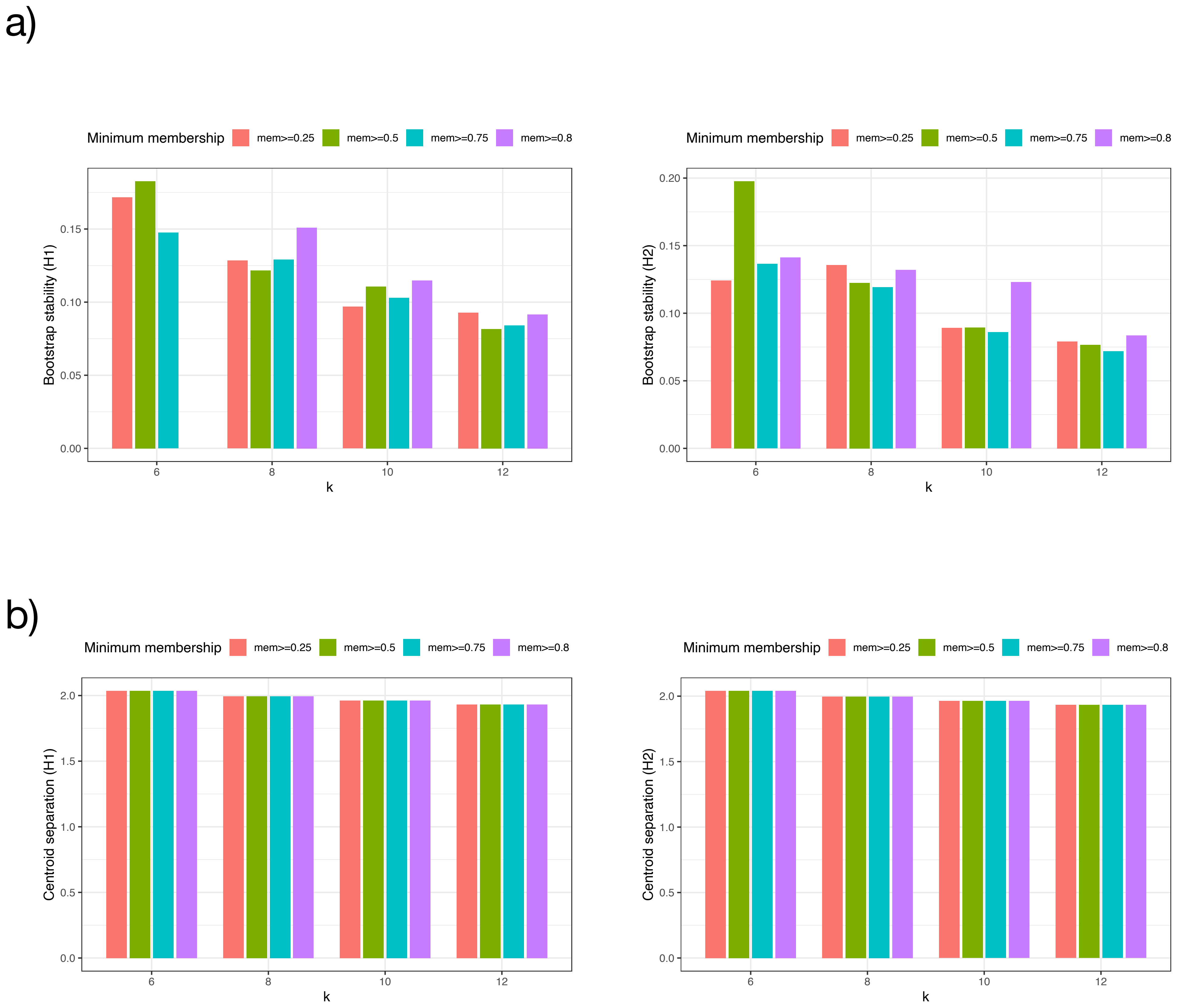


**Figure S4. Bootstrap stability and centroid separation across clustering parameters.** (a) Bootstrap stability of Mfuzz clusters for haplotypes H1 (left) and H2 (right) evaluated across different numbers of clusters (k = 6, 8, 10, 12) and minimum membership thresholds (mem ≥ 0.25, 0.5, 0.75, 0.8). Bars represent the proportion of genes consistently assigned to the same cluster across bootstrap iterations under each parameter setting. (b) Centroid separation for H1 (left) and H2 (right) clusters across the same values of k and minimum membership thresholds, measured as the average pairwise distance between cluster centroids.

a)

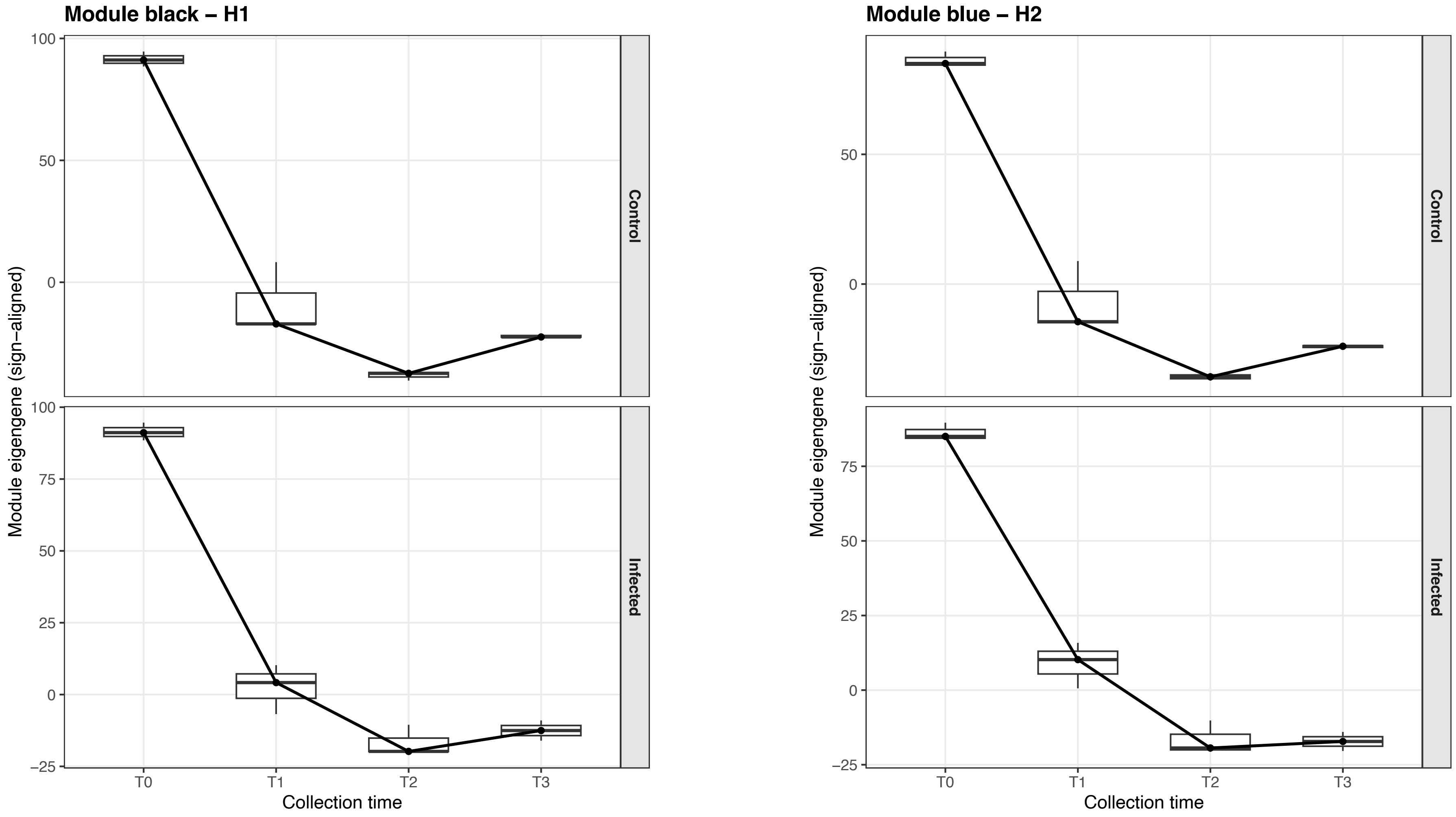


b)

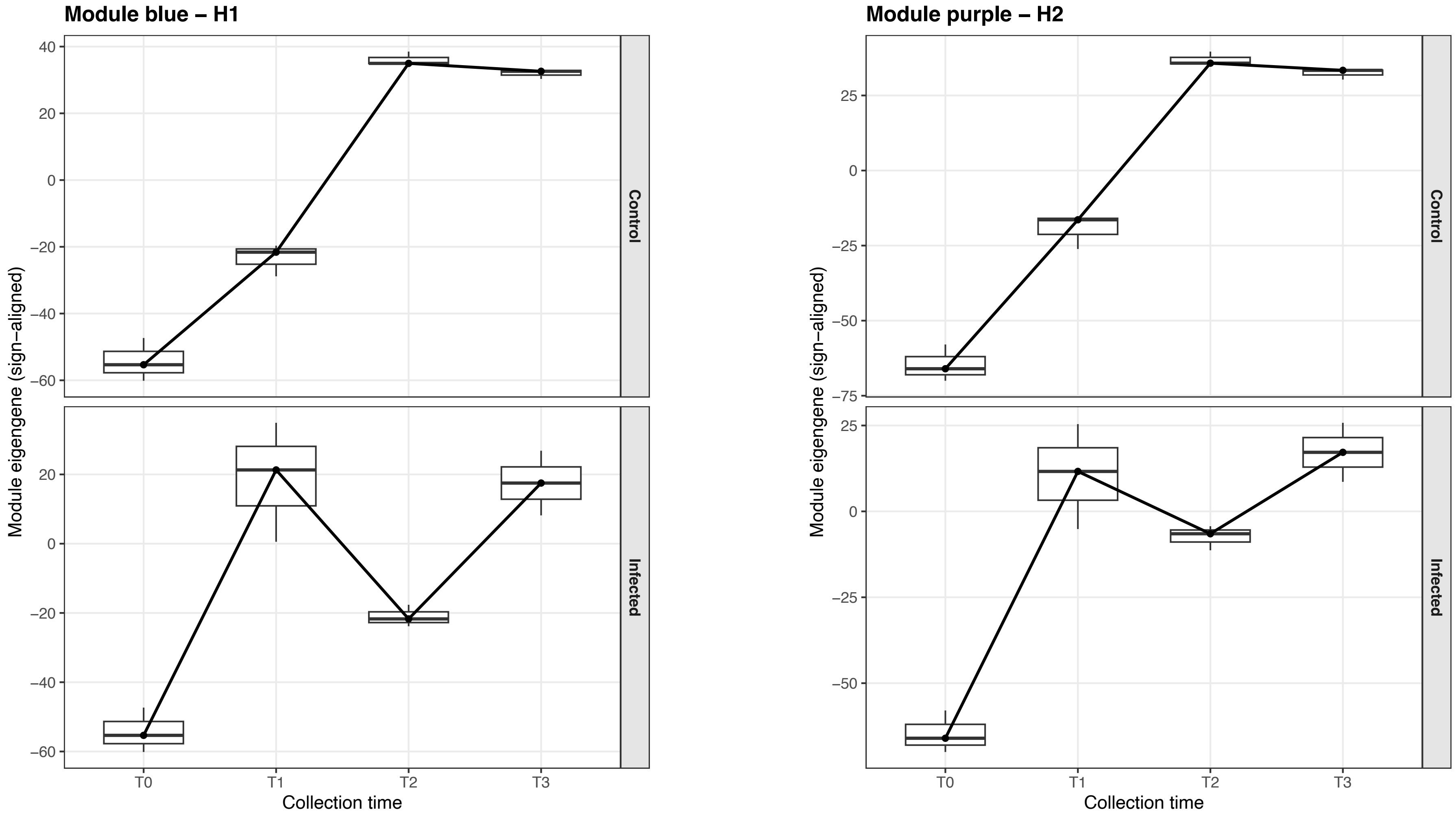


**Figure S5. Temporal eigengene trajectories of shared WGCNA modules across haplotypes.** (a) Time-course eigengene profiles for the H1–black and H2–blue modules. (b) Equivalent trajectories for the H1–blue and H2–purple modules. Boxplots reflect sample-level variation, while lines show mean eigengene trends across collection times (T0–T3).

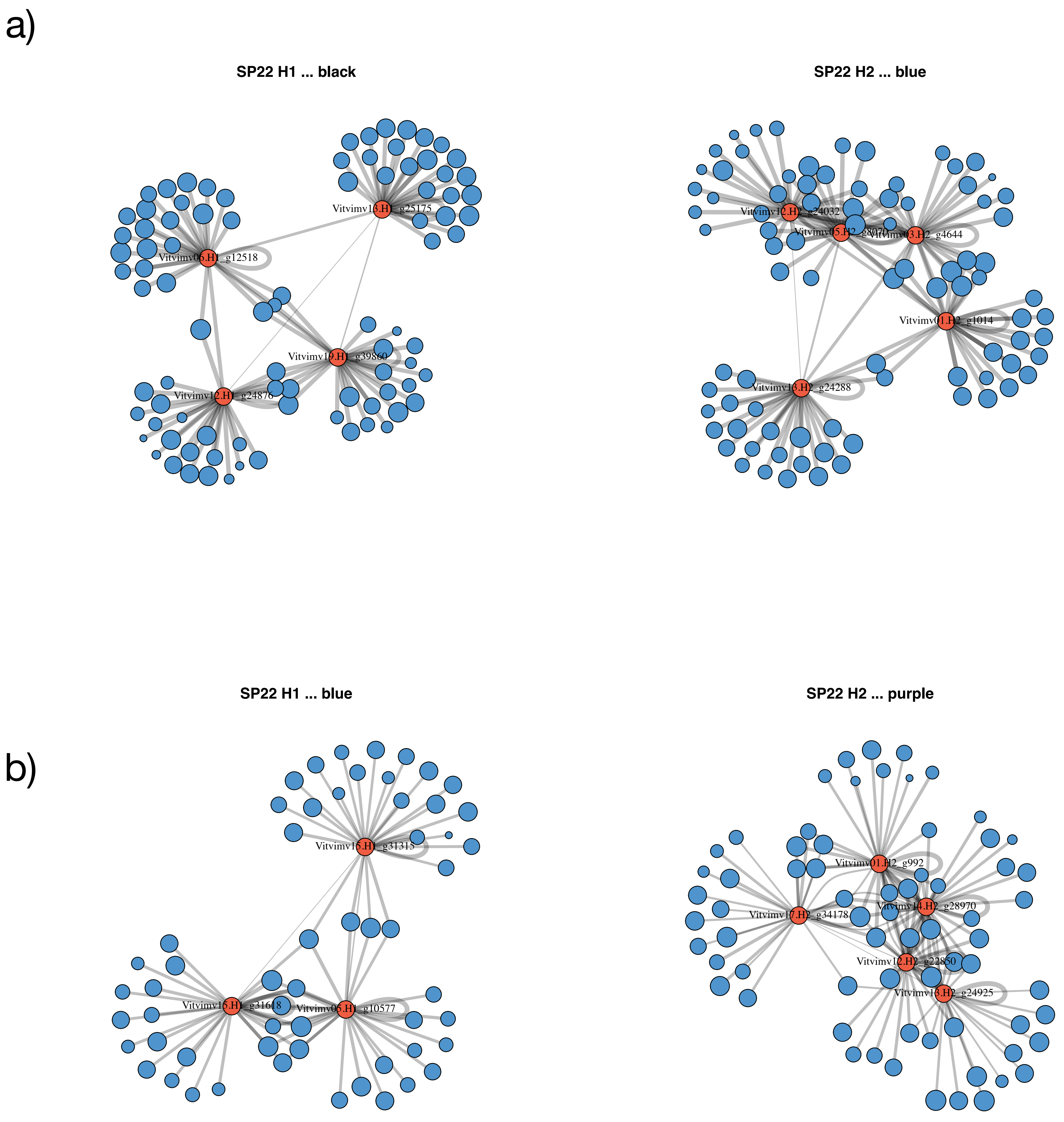


**Figure S6. Cross-haplotype WGCNA network modules sharing common genes.** (a) Subnetworks for modules containing shared genes between haplotypes: (a) H1–black vs H2–blue, and (b) H1–blue vs H2–purple. Hub genes are highlighted, with edge density reflecting intramodular connectivity.

a)

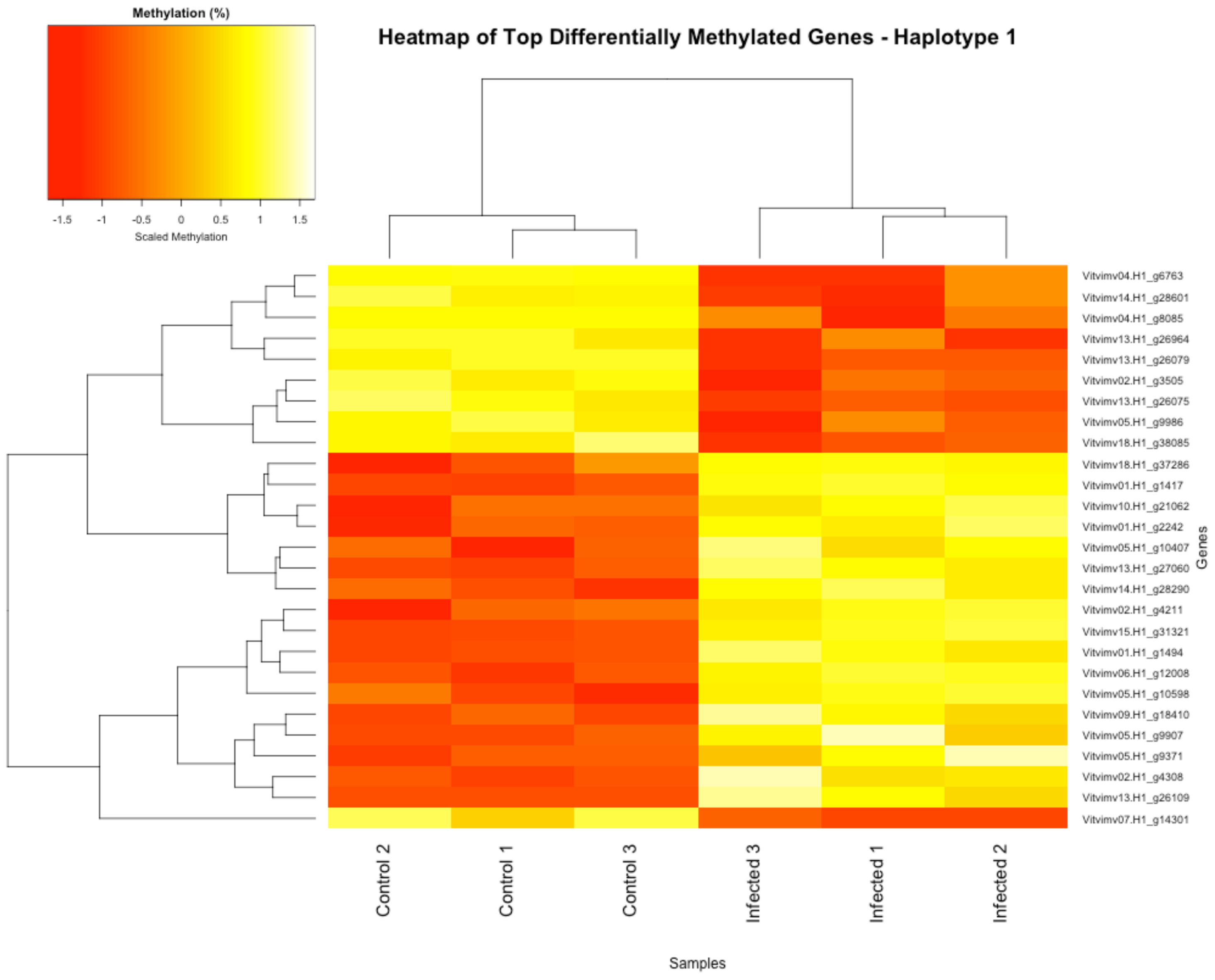


b)

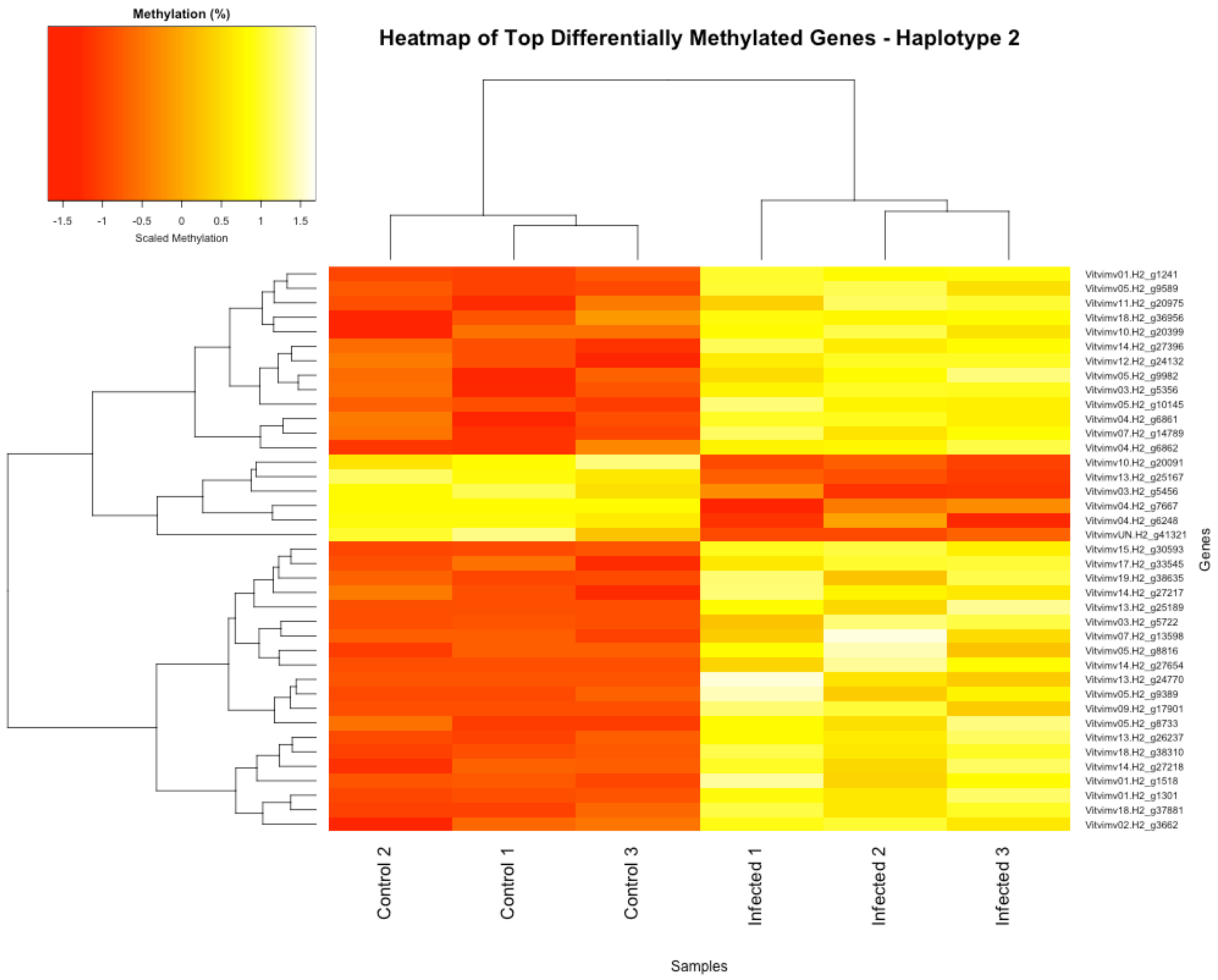


**Figure S7. Differentially methylated genes across haplotypes.** Heatmaps of the top differentially methylated genes (DMGs) in (a) Haplotype 1 and (b) Haplotype 2.
